# Adducin Regulates Sarcomere Disassembly During Cardiomyocyte Mitosis

**DOI:** 10.1101/2020.12.24.424022

**Authors:** Feng Xiao, Ping Wang, Shujuan Li, Suwannee Thet, Diana C. Canseco, Wataru Kimura, Xiang Luo, Ngoc Uyen Nhi Nguyen, Nicholas Lam, Waleed Elhelaly, Rohit Singh, Sakthivel Sadayappan, Mohammed Kanchwala, Chao Xing, Roger J. Hajjar, Joseph A. Hill, Hesham A. Sadek

## Abstract

Recent interest in understanding cardiomyocyte cell-cycle has been driven by potential therapeutic applications in cardiomyopathy. However, despite recent advances, cardiomyocyte mitosis remains a poorly understood process. For example, it is unclear how sarcomeres are disassembled during mitosis to allow abscission of daughter cardiomyocytes. Here we identify adducin as a regulator of sarcomere disassembly during mammalian cardiomyocyte mitosis. α/γ-adducins are selectively expressed in neonatal mitotic cardiomyocytes, and their levels decline precipitously thereafter. Cardiomyocyte-specific overexpression of various splice isoforms and phosphoforms of α-adducin in-vitro and in-vivo identified Thr445/Thr480 phosphorylation of a short isoform of α adducin as a potent inducer of neonatal cardiomyocyte sarcomere disassembly. Concomitant overexpression of this α-adducin variant along with γ-adducin resulted in stabilization of the adducin complex and persistent sarcomere disassembly in adult mice, which is mediated by interaction with α-actinin. These results highlight an important mechanism for coordination of cytoskeletal morphological changes during cardiomyocyte mitosis.

## Introduction

Cardiomyocyte loss is a leading cause of heart failure, which is a devastating progressive disease affecting over 30 million patients worldwide ^1^. Although limited cardiomyocyte renewal occurs in the adult mammalian heart, it is insufficient for restoration of contractile function following cardiomyocyte loss which results in cardiomyopathy ^2–7^ Therefore, stimulating cardiomyocyte regeneration is a major goal of heart repair in cardiomyopathy. Unlike some lower vertebrates which possess a life-long cardiac regeneration capacity ^8–14^, adult mammalian cardiomyocytes permanently withdraw from the cell cycle shortly after birth and have a very limited regenerative potential. Studies in neonatal mouse hearts showed that apical resection or myocardial infarction (MI) of postnatal day 1 (P1) mice leads to a robust regenerative response, however this capacity is lost by P7 which coinciding with post-natal cardiomyocyte cell cycle arrest ^15,16^ To date, several regulators of cardiomyocytes cell cycle have been identified^17–23^, however, it is not clear whether there are muscle-specific factors that regulate mitosis in cardiomyocytes.

The dense myofibrillar composition of cardiomyocytes poses a special challenge during mitosis, which is a consideration that does not exist in other non-striated cells. In an adult cardiomyocyte, sarcomeres occupy over 60% of cardiomyocyte cytoplasm ^24^, and anchor to the plasma membrane to induce deformation of the cell during contraction. An important and interesting feature of regenerating hearts is disassembly of cardiomyocyte sarcomeres ^15,25,26^, and their peripheral marginalization during cardiomyocyte mitosis^27^. This phenomenon has long been observed in dividing cardiomyocytes and has proven to be a reliable indicator of cardiomyocyte mitosis in the early postnatal heart ^15,17,22,28^ However, to date, the mechanisms that regulate sarcomere disassembly are not understood. Moreover, it is unclear whether regulation of sarcomere disassembly plays a role in loss of the regenerative capacity of the neonatal heart with advancing postnatal age. In the current study, we set out to identify regulators of cardiomyocyte sarcomere disassembly during mitosis. We found that the cytoskeletal regulatory protein adducin is a critical regulator of sarcomere disassembly.

## Results

### Adducin is predominantly expressed during the neonatal regenerative window and tightly associated with sarcomere disassembly

Immunostaining of the regenerating neonatal heart demonstrates clearance of cytoplasmic sarcomere and localization of troponin T (Tnnt2) to the periphery of cardiomyocytes during mitosis ^15^. This association of Tnnt2 with disassembled sarcomeres can thus be used to gain insights into other disassembly-associated proteins. Therefore we compared the Tnnt2-associated proteomic profiles by mass spectrometry (MS) after Co-Immunoprecipitation (Co-IP) with Tnnt2 antibody at two different developmental stages; 3 days after myocardial infarction (MI) at P1 (P1MI), a timepoint associated with robust cardiomyocyte proliferation, and 3 days after MI at P7 (P7MI), a time point which lacks mitotic induction of cardiomyocytes (Fig. 1B) ^15, 16^. Among 224 identified proteins, we found that 80% of them were cytoskeleton-associated proteins. Fig.1C lists selected cytoskeleton or sarcomeric proteins identified by MS. α-spectrin is one of the most abundant protein on the list of the P1MI sample but showed little expression in the P7MI sample. We also found that α- and γ-adducin are differentially associate with Tnnt2 during the regenerative window. Although both proteins were weakly detected by MS, they were only detected in the P1MI sample. Previous studies in non-muscle cells have shown that adducin regulates the interaction of spectrin with actin to form a lattice for mechanical support of plasma membranes As a downstream effector of several signaling pathways, adducin is known to be an important regulator controlling the interactions of spectrin, actin and other cytoskeleton and membrane proteins in erythrocytes ^30^, endothelial cells^31,32^, and the central nerve system^33^, however, almost nothing is known about its function in muscle cells or in cardiomyocytes. Adducin has three isoforms, α-adducin (ADD1), β-adducin (ADD2), and γ-adducin (ADD3), which are encoded by different genes but share highly homologous head, neck and tail domains^34^. Adducins form heterodimers through the interaction at the head domain of either α-/β- or α-/γ-adducin. α-/γ-Adducin is ubiquitously expressed in all tissues, while β-adducin is limited to erythrocyte or brain cells^35^. Pulldown of α-adducin by Tnnt2 was then confirmed with a reverse Co-IP in which Add1 antibody was used as bait and the resulting pulldown proteins were detected by Tnnt2 antibody (Fig. 1D). Association of adducin with sarcomeric protein α-actinin was also verified by Co-IP and WB (Fig. 1D). Examination of endogenous adducin protein expression by WB indicated that the expression of both α- and γ-adducin was downregulated with increasing postnatal age (Fig. 1E). We then examined the expression pattern of α-Adducin, γ-Adducin, α-spectrin, and α-filamin, which were 4 of the top hits from the MS screen, by immunohistochemistry (Fig. 1F). We found that adducin and spectrin only colocalized with disassembled sarcomere in proliferating cardiomyocytes, where they were associated with membranous and submembranous region. α-Filamin showed a prominent nuclear staining pattern in non-proliferation cells and a weak cytoplasmic pattern in proliferation cells. These results support the association of adducin with proliferating cardiomyocytes in the neonatal heart.

**Figure 1.**
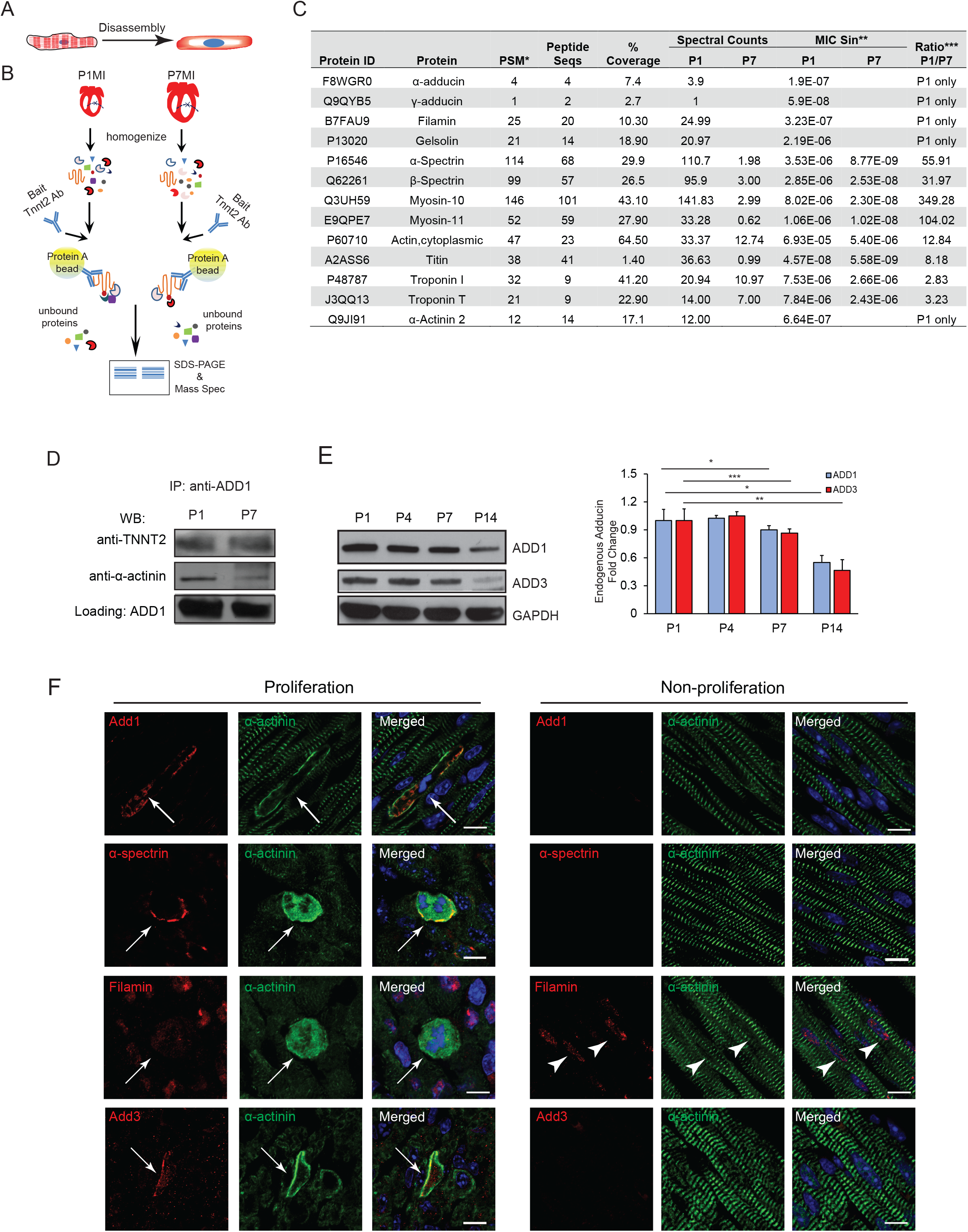
Identification of adducin as a key protein associated with sarcomere disassembly. **A.** Schematic shows the morphology of sarcomere disassembly. **B.** Schematic of Co-IP/MS by TNNT2 pulldown. **C.** Table lists selected TNNT2 associated proteins identified from the Co-IP/MS. * PSM: Peptide Spectrum Matches. Number of spectra assigned to peptides that contributed to the inference of the protein. **MIC Sin: The normalized spectral index statistic for the protein for the specific group. Calculated from the intensity of fragment ions in each spectrum assigned to a protein. ***Ratio: Quantitative ratio for protein between groups derived from the MIC Sin value. **D.** Coimmunoprecipitation of ADD1 from total heart extracts at P1 and P7, probed for TNNT2 and α-actinin. **E.** (Left) Representative WB to show endogenous expression profile of ADD1 and ADD3 at different ages. (Right) Quantification of adducin expression level relative to internal control GAPDH. (n=3 per group). **F.** Expression of four selected cytoskeletal proteins from MS results, α-, γ-adducin, spectrin, and filamin were stained in neonatal hearts, showing profile of proliferation (left) and non-proliferation (right) cardiomyocytes. Arrows point to representative cells. Arrow heads in filamin staining images refer to nucleus location. Scale bar=10 μm.

### Overexpression of α-adducin isoform 2 can disassemble sarcomeres in juvenile

Alternative splicing is an important post-transcriptional regulation in muscle-specific genes^36,37^. It also plays an important role in heart development and cardiomyopathy^38–40^. So far nearly all publications about α-adducin were focused on variant 1 which encodes a full length Add1. According to information from Ensembl, Add1 has 15 splice variants which results in two major isoforms, isoform 1 and 2. Their difference lies in the expression of exon 15, which has an in-frame stop codon (Fig. 2A Top). Therefore, exon 15 inclusion will transcribe variant 2 which encodes a shorter isoform 2 lacking ~100 amino acids at the C-terminus. On the other hand, transcription with exon 15 exclusion will make a full-length isoform 1 of α-adducin (Fig. 2A Bottom). We examined endogenous expression level of isoform 2 in heart at different age by WB (Fig. 2B). We found that the expression levels of isoform 2 followed a similar pattern as isoform 1 where it also declines with postnatal age in the mouse heart. To elucidate a potential role for Add1 in sarcomere disassembly in cardiomyocyte, we generated constructs to express WT isoform 1 (Add1 i1) and 2 (Add1 i2) individually, and we packaged them into Adeno-associated virus (AAV). We used cardiotropic AAV6 carrying the different isoforms in neonatal rat ventricle myocytes (NRVM) (Fig. 2C) to determine the effect of adducin expression on sarcomere disassembly. AAV6-GFP was included as a control. Purified viruses were used to infect cultured NRVM at a titer of 5 x 10^4^ vector genome per cell (vg/cell) for 72 hrs prior to fixation. To quantify the extent of sarcomere disassembly, we pre-defined three types of patterns; assembled, partial disassembled, and fully disassembled sarcomeres. Partial disassembled sarcomere refers to myocytes with any visible sarcomeres, while fully disassembled sarcomere are myocytes without any sarcomeres. We found that 61.9% of NRVMs with Add1 i1 overexpression showed partial disassembled sarcomeres (Fig. 2C-i), while 10.8% of Add1 i1 positive cells show completely disassembled sarcomeres where no sarcomeres were detected (Fig. 2C-ii). Overexpression with Add1 i2 resulted in 49.1% partial disassembly and 50.8% complete disassembly (Fig. 2C iii, iv). These results indicate that Add1 i2 overexpression is more effective in inducing sarcomere disassembly than Add1 i1. We then sought to generate a cardiac-specific transgenic (TG) model where Add1 i2 expression is driven by an α-MHC promoter (Fig. 2D Top). Add1i2 transgenic mice did not show a significant difference in cardiac size and morphology by H&E staining (Fig. 2D Bottom left), or HW/BW (Fig. 2D Bottom right). When we compared adducin overexpression pattern at P14 and P28, we noted a relative decrease in Add1 at P28 compared to P14 hearts (Fig. 2E). Importantly, we found that P14 hearts showed evidence of sarcomere disassembly, while P28 hearts did not (Fig. 2F). Assessment of left ventricular systolic function by echocardiography did not show any appreciable difference between WT and Add1i2 TG hearts (fig. S1A). Finally, we observed a significant increase in pH3-positive cardiomyocytes Add1i2 TG at p14 (fig. S1B), with no change in cardiomyocyte size (fig. S1C). These results suggest that cardiomyocyte-specific overexpression Add1 i2 causes a transient increase in sarcomere disassembly, which does not persist to adulthood.

**Figure 2.**
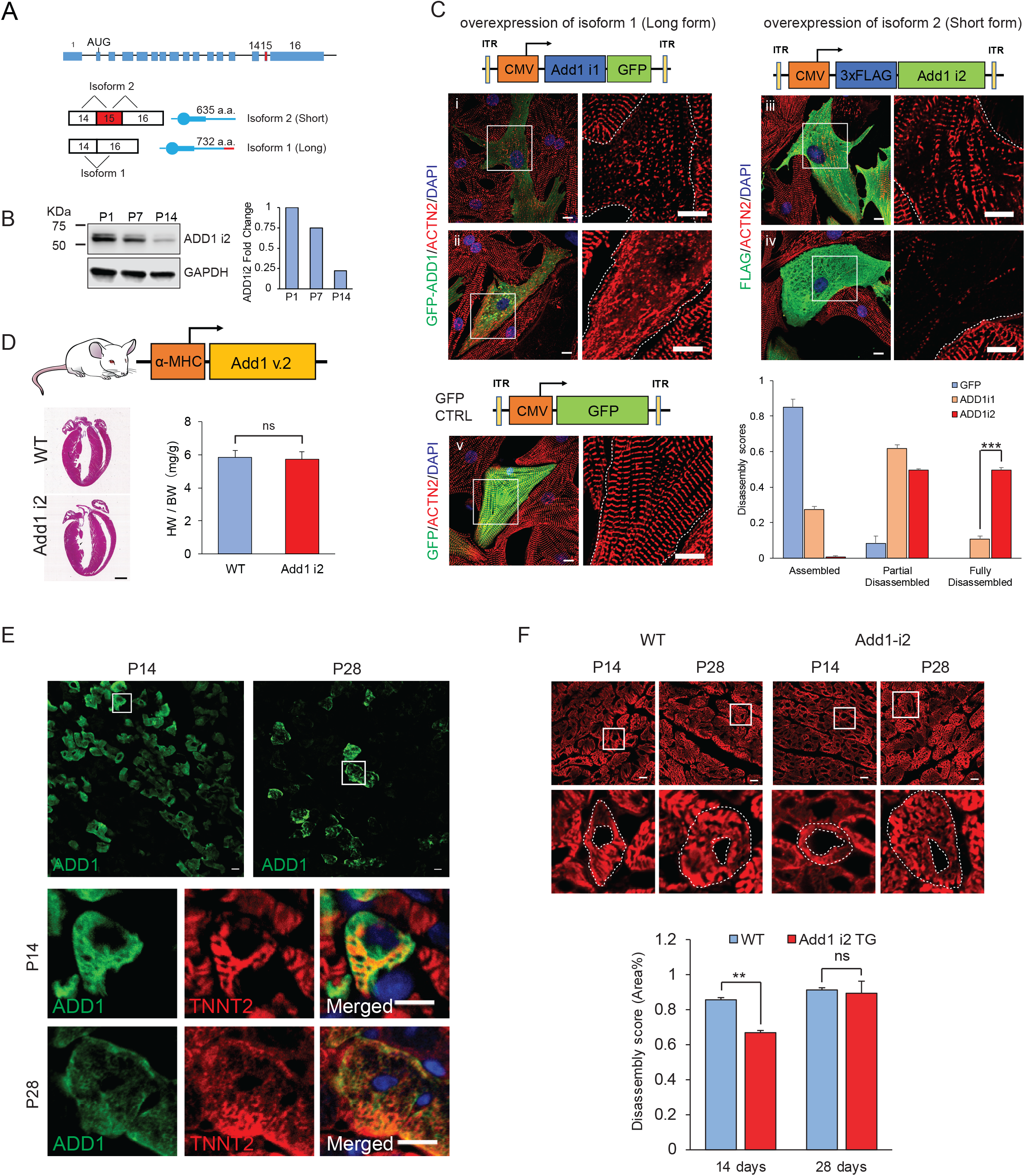
Overexpression of Add1 isoform 2 disassembles cardiomyocyte sarcomeres. **A.** (Top) Schematic of exons and introns of *Add1* gene. Red rectangle represents exon 15 which is the target of alternative splicing. (Bottom) Schematic shows inclusion or exclusion of exon 15 in mRNA alternative splicing events. The corresponding translated protein schematic is shown next to the mRNA. Inclusion of exon 15 has an in frame stop code which generates ADD1 isoform 2, composing 636 amino acids (a.a.). Exclusion of exon 15 translates into ADD1 isoform 1, composing 732 a.a. **B.** Western blot shows endogenous profile of ADD1 isoform 2 at different ages. GAPDH is used as internal control. **C.** Representative images of NRVM transduced with AAV6-Add1i1 (i, ii), AAV6-Add1i2 (iii, iv), and AAV6-GFP (v) at day 3. Schematics of AAV expression cassettes are shown with each group images. Cells were fixed and immunostained with α-actinin (ACTN2, red) to reveal sarcomere organization. Green fluorescence represents either Adducin isoform 1 and 2, or GFP. High magnification images of squared regions are shown on the right. Dotted lines indicated the boarder of cardiomyocytes. (i) and (iii) represents partial disassembled sarcomeres. (ii) and (iv) represents fully disassembled sarcomeres. (v) represents assembled sarcomeres. Percentage of sarcomere assembly changes by AAV are shown in the graph. **D.** (Top) Schematic to show the strategy of generating Add1 isoform 2 single transgenic line. (Bottom left) H&E staining of WT and cardiac-specific Add1 i2 transgenic. Scale bar = 1 mm. (Bottom right) Heart weight to body weight ratio in control and cardiac-specific Add1i2 transgenic. **E.** (Top) Images to show expression pattern of ADD1 i2 transgenic at P14 and P28. (Bottom) High magnification images of squared region from the top panels. Scale bar = 10 μm. **F.** (Top) Representative images to show sarcomere pattern from ADD1 i2 transgenic at P14 and P28. Insets are high magnification images of single cardiomyocyte (squared regions). Dotted lines are drawn around outer and inner borders of sarcomeres. (Bottom) Quantification graph to show the extent of sarcomere disassembly. Disassembly scores were defined as: (Outer Area – Inner Area) / Outer Area. Age matched litter maters of WT mice were used as controls. At P14, ADD1 i2 transgenic mice have significantly thinner sarcomeric zone with central clearance compared to controls. Data are presented as mean ± s.e.m. *P<0.05, **P<0.01, ***P<0.001; statistical significance was calculated using a two-tailed *t*-test.

### Role of α-adducin phosphorylation in neonatal cardiomyocyte mitosis

Several adducin phospho-sites that regulate differential expression patterns have been previously identified. To understand whether adducin phosphorylation is involved in regulation of cardiomyocyte sarcomere disassembly, we examined the endogenous expression pattern of phospho-isoforms of α-adducin in the neonatal mouse heart (Fig. 3A). Previous studies have shown that phosphorylation of α-adducin at Thr445 and Thr480 in the neck domain enhances the recruitment of spectrin to F-actin ^31^. Staining of neonatal hearts with anti-phospho-Thr455 antibody showed localization to the cellular cortex only in myocytes with disassembled sarcomere (Arrows), but in the nucleus in nonproliferation cardiomyocytes (Arrow heads). It has also been reported that α-adducin is phosphorylated by cyclin-dependent kinase 1 (CDK1) at Ser12 and Ser355 in the head domain to associate with mitotic spindles ^32^. We found that mitotic cardiomyocytes demonstrated colocalization of Ser355 in the cytoplasm of mitotic cardiomyocytes (Arrow). Other studies have reported that Ser716 and Ser726 (S714 and S724 in mouse respectively) are located in a C-terminal myristoylated alanine-rich C kinase substrate-related domain (MARCKS), and are substrates for protein kinases A (PKA) and C (PKC). Phosphorylation of this domain inhibits recruitment of spectrin to actin filaments and induces disassembly of spectrin-F-actin meshwork^41^. Our results indicate that S724 phosphorylation does not occur in neonatal cardiomyocytes in vivo. Additionally, adducin can be phosphorylated by PKA at Ser408, Ser436, and Ser481 which inhibits its binding to spectrin-F-actin complex^42^. We found that phosphorylation of S724 or S481 did not occur in neonatal cardiomyocytes. We further examined the correlation of phospho-adducin expression with cardiomyocyte cell cycle stage both *in vitro* and *in vivo.* Immunofluorescent staining with pT445 antibody in NRVM (Fig. 3B) and P4 heart (fig. S2A) both confirmed the correlation of adducin expression with cardiomyocyte mitosis. In prophase pT445 appears to be nuclear, while in prometaphase with the initiation of disassembly adducin pT445 is expressed both in the cytoplasm and the mitotic chromosome. Adducin expression remains cytoplasmic through telophase. WB of adducin phospho-T445 expression pattern indicated that its expression decreases with age, which follows the same pattern of WT Add1 (Fig. 3C). This expression pattern of phospho-adducin T445/T480 during cardiomyocyte mitosis suggests that phosphorylation on T445 and T480 might play an important role in sarcomere disassembly.

**Figure 3.**
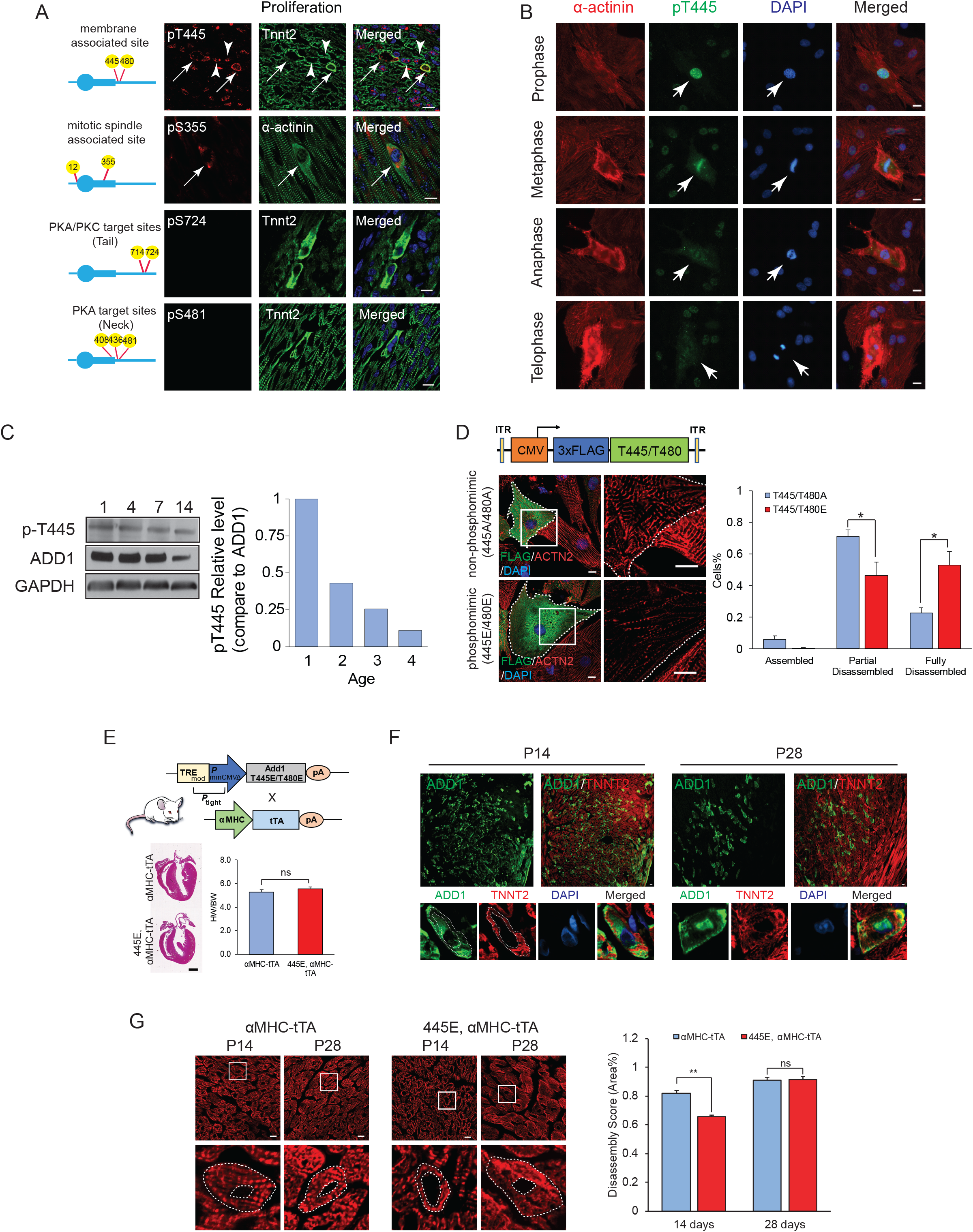
Role of Add1 i1 phospho-mimic on sarcomere disassembly. **A.** Endogenous phospho-adducin expression pattern in neonatal mouse heart. Schematics of phosphosite locations is shown on the left. Right: immunostaining images show co-staining images of anti-phospho-site antibody with sarcomeric protein TNNT2 or α-actinin. From top to bottom are Phospho-sites T445/T480, S12/S355, S714/S724, and S408/S436/S481. Arrows indicate phospho-adducin colocalized with disassembled sarcomeres. Arrow heads indicate colocalization with assembled sarcomeres. **B.** Endogenous expression of phospho-adducin T445 in NRVM as stained with α-actinin (red), anti-phospho-adducin pT445 (green) and DAPI. Phospho-adducin translocates from the nucleus to the cytoplasm during mitosis. **C.** (Top): WB examination of endogenous phospho-T445 expression level. (Bottom) Quantification of pT445 expression level in relative to ADD1 expression level. **D.** Overexpression of adducin phospho- or non-phospho mimic T445/T480 in NRVM as induced by AAV6. (Top left) Schematic diagram of AAV shuttle vector shows that adducin mutants are expressed with 3 tandem FLAG epitopes in frame at the N-terminus. (Bottom left) Immunofluorescent staining with α-actinin antibody to show sarcomeric structure. Adducin mutants are detected by Flag antibody in green. High magnification images of representative region (white box) were shown on the right. Dotted lines were drawn around the cell boarder. (Right) Quantitative analysis of disassembled sarcomeres induced by adducin overexpression. Scale Bar = 10 μm. **E.** (Top) Schematics to show strategy of generating cardiac specific inducible Add1 i1 phospho single transgenic (p-TRE T445E/T480E, α-MHC tTA). (Bottom left) H&E staining of control and cardiac-specific Add1 i1 phospho transgenic. Scale bar = 1 mm. (Bottom right) Heart weight to body weight ratio in control and cardiac-specific phospho-single transgenic. **F.** (Top) Images to show expression pattern of phospho-Add1i2 transgenic at P14 and P28. (Bottom) High magnification images of selective phospho-adducin mimic overexpression cardiomyocytes. At P14, we saw phospho-mimic expresses both in cytoplasm and on membrane (Bottom left). At P28, phospho-mimic adducin expresses on membrane and in nucleus (Bottom right). Scale bar = 10 μm. **G.** (Left) Representative images to show sarcomere pattern from phospho-ADD1 i1 transgenic at P14 and P28. *Insets* are high magnification images of single cardiomyocyte. Dotted lines are drawn around outer and inner edges of cardiomyocytes. (Right) Quantitative graph to show sarcomere disassembly. Disassembly scores were defined the same way as that of Add1i2 transgenic in Fig. 2F. Scale bar = 10 μm. Scale bar= 10 μm. Data are presented as mean ± s.e.m. *P<0.05, **P<0.01, ***P<0.001; statistical significance was calculated using a two-tailed *t*-test. Data are presented as mean ± s.e.m. *P<0.05, **P<0.01, ***P<0.001; statistical significance was calculated using a two-tailed *t*-test.

To determine whether overexpression of phosphorylated adducin in cardiomyocyte influences sarcomere disassembly, we constructed a series of adducin mutants to mimic the phosphorylated and non-phosphorylated adducin by site-directed mutagenesis (phospho-mimic and phospho-silent). Substitution with Glu mimics the adducin phosphorylation form, while mutation to Ala mimics the non-phosphorylated form. The constructs were further packaged with helper plasmid to generate a recombinant AAV6 virus. The sarcomere morphology was examined by staining with α-actinin antibody and colocalizing with FLAG positive cells. Our results demonstrate that nearly 60% Add1 positive cardiomyocytes with phospho-mimic overexpression (T445E/T480E) demonstrate full sarcomere disassembly compared to cardiomyocytes infected with the phospho-silent (T445A/480A) form, which mainly induced partial sarcomere disassembly (Fig. 3D). We also generated phospho-mimic and phospho-silent mutants of the phospho-sites S12/S355 and S714/S724. Infection of NRVMs with AAV6 carrying these phospho-mutants did not show a significant increase in complete sarcomere disassembly (fig. S2B and S2C), although the phospho-silent form of S714/S724 did show higher rates of complete disasembly compared to the phospho-mimic form. Therefore, we were focused on phospho-sites T445/T480 to study their role in sarcomere disassembly.

To determine whether forced expression of the constitutively active membrane-associated form of *Add1* can induce sarcomere disassembly *in vivo,* we generated an inducible pTRE-T445E/T480E transgenic (TG) mouse line, where the phospho-isoform of adducin is selectively expressed in cardiomyocytes only after crossing with αMHC-tTA mouse and withdrawal of doxycycline (Tet-off) (Fig. 3E). The transgenic didn’t show significant change on heart size or HW/BW at P28 (Figure 3E). Similar to the aforementioned Add1 i2 single transgenic line, we found that the phospho-adducin positive cardiomyocytes had a patchy pattern of expression, and we observed decreased expression from P14 to P28 (Fig. 3F, Top). Further examination of the sarcomeric pattern in the phospho-mimic positive cardiomyocytes showed cytoplasmic expression of adducin with sarcomere disassembly at P14, which did not persist until P28 cardiomyocytes, which showed normal sarcomere structures (Fig. 3F bottom). Assessment of the disassembly score confirmed that while cardiomyocytes were disassembled at P14, they had normal morphology by P28 (Fig. 3G), which is reminiscent of the pattern we noted earlier with the Add1i2 TG (Fig. 2F). This suggested that overexpression of Add1 i2 or phospho-mimic Add1 i1 help to extend the window of sarcomere disassembly but fail to maintain this pattern until adulthood. Although we detected pH3 signals in hearts at P14, there is no significant difference compared with that of the control (fig. S3A). The cardiomyocyte cell size also shows no difference either at P14 (fig. S3B).

### Concomitant expression of α and γ-adducins is required for protein stabilization

Previous reports have mostly focused on α-adducin primarily because it is thought to be the limiting factor in a heterodimer formation with either γ or β adducin, or even a homodimer with itself ^43^. Support of this notion comes from elegant studies in Add1 KO models which suggested that α-adducin deletion resulted in a greatly reduced expression of γ-adducin. However, in our studies, failure to disassemble sarcomere in both single Add1 TG after P14 raises an important question: Is γ-adducin needed for α-adducin function in vivo? In our studies, endogenous expression profile of adducins showed that γ-adducin expression becomes negligible at P14 (Fig. 1E). This might suggest that Add1 single TG lines lose the ability to disassemble sarcomeres beyond P14 secondary to the age-dependent loss of Add3. To evaluate the stability of Add1 in relation to Add3, we performed *in vitro* transfection with Add1 var.1 and var. 2 in the presence or absence of Add3. To avoid adducin antibody cross reaction between α-adducin and γ-adducin, we transfected cells with Add1 variants with 3 tandem FLAG tag at N-terminus and/or plasmids of Add3 with 3 tandem TY1 tag at N-terminus. 24 hours after transfection, the cells were treated with 80 μM cycloheximide (CHX) to inhibit protein translation. We collected cells at multiple time points after treatment and examined protein levels by WB. Intriguingly, we found that that both α-adducin i1 and i2 became unstable upon CHX treatment in the absence of γ-adducin. However, in the presence of γ-adducin, the stability of α-adducin isoforms 1 and 2 were greatly enhanced. (Fig. 4A). This result indicates that the failure of the single transgenic lines to induce pronounced disassembly beyond P14 might be related to the absence of endogenous γ-adducin at later timepoints.

**Figure 4.**
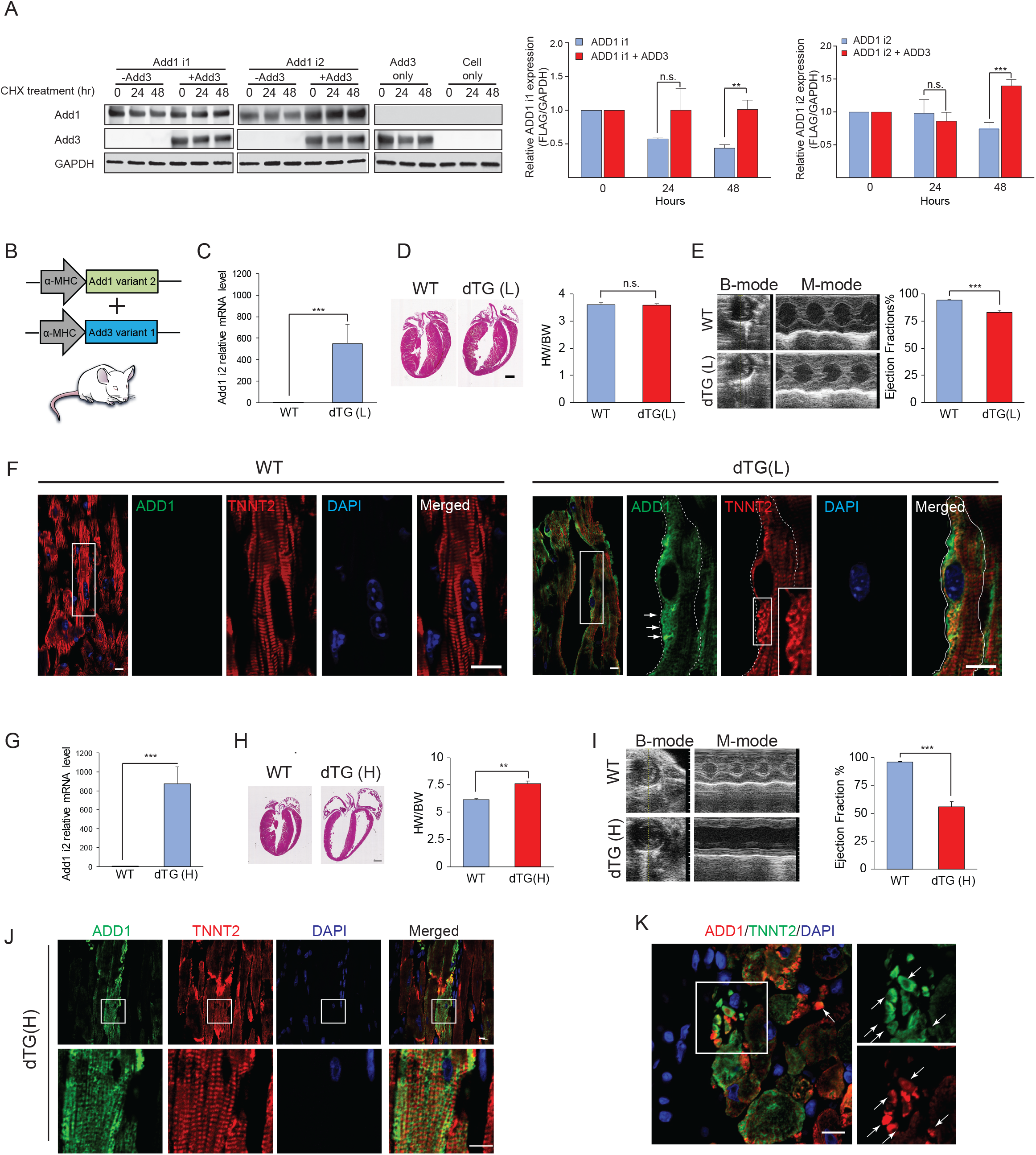
Co-expressing Add1 isoform 2 together with Add3 enhance adducin stability. **A.** HEK293T cells were transfection of variant 1(p-3xFlag-Add1i1) and Add1 variant 2 (p-3xFlag-Add1i2) in the presence or absence of Add3 (p-3xTy1-Add3). 24 hours after the transfection, the cells were treated with 80 μM cycloheximide (CHX) to inhibit protein translation. Cells were collected 0, 24, and 48 hours after treatment. (Left) WB to show protein level. (Right) Measurement of relative ADD1i1 or Add1i2 protein level (calculated as FLAG vs GAPDH) in the absence or presence of ADD3. Data are presented as mean ± s.e.m. *P<0.05, **P<0.01, ***P<0.001; statistical significance was calculated using a one-tailed *t*-test. **B.** Schematic diagram to show the constructs of generating Add1 i2 and Add3 double transgenic. Both expression constructs were driven under an α-MHC promoter. **C.** QPCR results to show the mRNA level of Add1i2 in 2-month-old double transgenic with low expression. **D.** (Left) H&E staining of control and cardiac-specific Add1 i2/Add3 double transgenic with low expression. Samples are of two months old mice. Scale bar = 1 mm. (Right) Heart weight to body weight ratio in WT and low expression double transgenic. **E.** Echocardiography (Left) and left ventricular systolic function quantification by ejection fraction (Right) in 2-month old ADD1 i2/ADD3 double transgenic (Low expression). n=3 for each group. **F.** Longitudinal views to show sarcomere pattern from 2 months old WT and Adducin double transgenic (Low expression). The high magnification images of boxed region in low magnification image are shown on the right. Dotted lines the border where adducin is expressed. *Inset* shows partial disassembled sarcomeres (disorganized but not absent sarcomeres). The corresponding adducin expression are indicated by arrows. Scale bar= 10 μm. **G.** QPCR results to show the mRNA level of Add1i2 in 2-month-old double transgenic with high expression. **H.** (Left) H&E staining of control and cardiac-specific Add1 i2/Add3 double transgenic with low expression. Samples are of 2-month-old mice. Scale bar = 1 mm. (Right) Heart weight to body weight ratio in WT and high expression double transgenic. **I.** Echocardiography (Left) and left ventricular systolic function quantification by ejection fraction (Right) in 1-month-old ADD1 i2/ADD3 double transgenic (high expression). (n=3 for each group). **J.** (Top) Representative images to show sarcomere pattern from 1-months old ADD1 i2/ADD3 double transgenic (high expression). (Bottom) Enlarged images of *Insets* from above panels. **K.** (Left) ADD1 i2/ADD3 double transgenic with high expression has evidence of cardiomyocytes fragmentation. (Right) High magnification of boxed region on the left. Arrows point to sarcomere fragments where ADD1 and TNNT2 are colocalized. Scale bar = 10 μm. Scale bar= 10 μm. Data are presented as mean ± s.e.m. *P<0.05, **P<0.01, ***P<0.001; statistical significance was calculated using a two-tailed *t*-test.

To test this hypothesis, we generated a cardiac specific double transgenic (dTG) mouse line co-expressing Add1 variant 2 gene and Add3 gene. This line was generated by injecting two independent expression cassettes together (Fig. 4B). Out of 35 founders in total, 11 were positive dTG. Immunostaining by Add1 and Add3 confirmed that both α- and γ-adducin are expressed and colocalized in cardiomyocytes (fig.S4A). Based on qPCR results of Add1 i2 expression levels, we separated lines into two groups based on the level of expression (Fig. 4C). It is important to note here that endogenous Add1i2 expression levels at baseline are not detectable after P14, which is why the transgenic lines show high level relative mRNA expression in comparison.

We first examined the lines with lower expression levels (dTG-L). We found that cardiac morphology and HW/BW at 2-months of age were similar to WT (Fig. 4D - representative lower expression line). However, LV systolic function of dTG-L mice was slightly lower than WTs (Fig. 4E). Intriguingly, we found that sarcomeric structures showed partial disassembly in cardiomyocytes were adducin was overexpressed (Fig. 4F). This morphology appeared as disorganized sarcomeres without clear striations (Fig. 4F). However, we did not observe the classic pattern of sarcomere disassembly with sarcomeric clearance. Next we examined the phenotype of the high adducin expression group (dTG-H). The mRNA level of high expression group dTG-H was about 1.6 times higher than the low expression group (Fig. 4G-representative higher expression line). Heart sections of dTG-H at two month of age showed ventricular dilation (Fig. 4H, left), and an increased HW/BW ratio (Fig. 4H, right). Echocardiography showed a markedly depressed LV systolic function with a left ventricular ejection fraction (LVEF) of 56% (Fig. 4I). We observed some partial disassembly with focal localization of TNNT2 to the intercalated disk (Fig. 4J), although some myocytes did not demonstrate significant sarcomere disorganization (Fig. 4J lower row). We also observed cardiomyocyte disruption and fragmentation suggesting increased cell death (Fig. 4K). which was further confirmed by TdT-mediated dUTP nick end labeling (TUNEL) assay (fig. S4B). Next we assessed cardiomyocyte size by WGA staining, and we found that the dTG-H had significantly high percentage of small cardiomyocytes, a lower percentage of intermediate size cardiomyocytes, and a similar percentage of large cardiomyocytes (fig.S4C). These results indicate that concomitant expression of Add3 and Add1 stabilizes the adducin complex, and that both dTG-L and dTG-H display partial sarcomere disassembly, with dTG-H showing evidence of widespread cardiomyocyte death and depressed LV function.

### Generation of Add1/Add3 Phospho-adducin Double Transgenic Mouse Model

As outlined earlier, the phospho-single TG T445E/T480E displays sarcomere disassembly in the neonatal but not the adult heart (Fig. 3G), likely due to lack of Add3 expression in the adult heart. Given the enhanced disassembly of neonatal cardiomyocytes with forced phospho T445E/T480E expression, we integrated the design of phoshpho-mimic transgenic with Add1i2/Add3 dTG to generate a new cardiac-specific double transgenic which co-expresses Add1 i2 phospho mimic T445E/T480E and WT Add3 (p-dTG) (Fig. 5A top). H&E staining of hearts from these mice demonstrated that the adult p-dTG hearts are bigger than the WT (Fig. 5A bottom), which was confirmed by HW/BW measurement (Fig. 5B). LVEF was mildly depressed in two-month old mice, although it was not significantly changed in younger mice (Fig. 5C). Examining adducin mRNA level by qPCR confirmed the increased level of expression of pAdd1i2 compared to WT hears (Fig. 5D). Immunostaining of Add1 and Add3 confirmed that both α- and γ-adducin are expressed and colocalized in the cytoplasm of cardiomyocytes (fig. S5A). We then examined the sarcomeric structure by troponin staining at P7, P14, P21 and 2 months. The results demonstrate that p-dTG cardiomyocytes display disassembled sarcomeres which was evident at all timepoints (Fig. 5E I,ii, iii & iv), although the disassembly appears more pronounced in early neonatal stages. In fact, most adult cardiomyocytes displayed partial disassembly that appeared to be localized a zone surrounding the nuclei and in the periphery of cardiomyocytes with complete clearance of the sarcomeres in these zones. Examination of cardiomyocyte mitotic markers of these hearts demonstrated an increase in pH3 and aurora B kinase-positive cardiomyocytes both at P21 and 2 months (fig. S5B & C), with no change in cell size (fig. S5D), or nucleation (fig. S5E).

**Figure 5.**
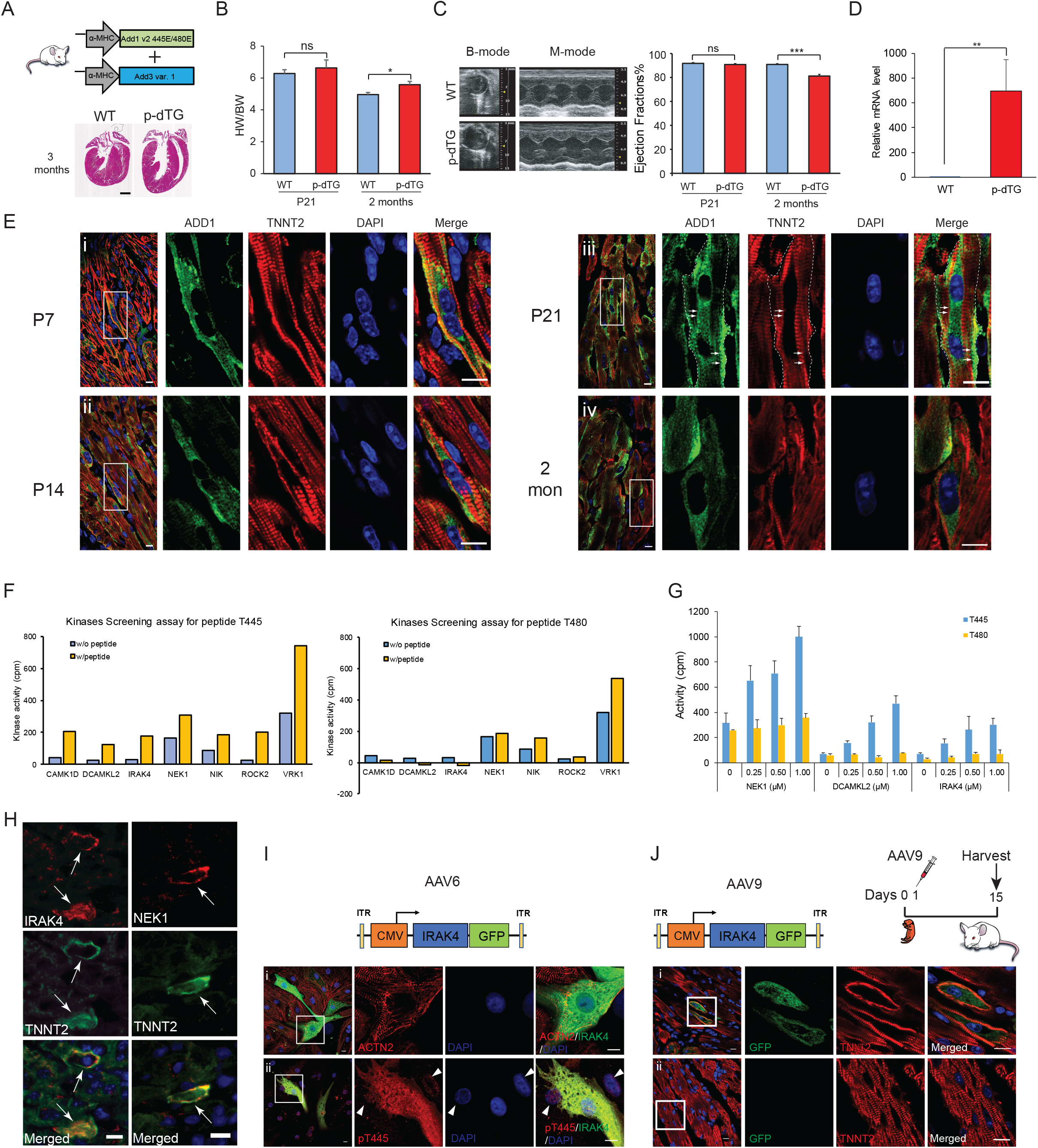
Phospho-mimic double transgenic promotes sarcomere disassembly. **A.** (Top) Schematic diagram of the constructs of generating phospho-mimic Add1 i2 and Add3 double transgenic. Glu substitution on T445 and T480 mimic phosphorylation form of Add1 i2. Both expression constructs were driven under an α-MHC promoter. (Bottom) H&E staining of control and cardiac-specific phospho-Add1 i2/Add3 double transgenic. Samples are of two-month-old mice. Scale bar = 1 mm. **B.** Heart weight to body weight ratio in WT and phospho double transgenic at P21 days and 2 months. (n=3 for P21, and n=5 for 2 months). **C.** Echocardiography (Left) and left ventricular systolic function quantification by ejection fraction (Right) in P21 and 2 months old ADD1 i2/ADD3 double transgenic (Low expression). (n=3 for P21, and n=5 for 2 months). **D.** QPCR of mRNA level of Add1i2 in 2-month-old phospho-double transgenic. **E.** Sarcomere pattern of phospho-double transgenic at P7 (i), P14 (ii), P21 (iii) and 2-month-old (iv). The high magnification images of boxed region are shown on the right side of zoom out image. Dotted lines circle out the boarder where adducin expresses outside sarcomeres. Arrows refer to regions with high level of adducin expression. Note the clearance of sarcomeres where adducin is expressed. Scale bar= 10 μm. **F.** Large scale kinase screening assay identified 7 candidates that potentially phosphoorylate T445 (left) and T480 (right) sites. Corrected kinase activity values (raw value minus sample peptide background, in cpm) with sample peptide at one concentration in 245 Ser/Thr peptide kinase assays. Blue: Kinase with sample peptide; yellow: Kinase activity w/o sample peptide. **G.** Serine/Threonine Kinases Hit Assay to verify 3 kinases, NEK1, DCAM1 and IRAK4, selected from initial kinase screening assay. Each kinase was tested at three concentrations in triplicates. These results suggested that the kinases NEK1, DCAMKL2, IRAK4 all show a concentration-dependent ability to phosphorylate the sample peptide “T445” and “T480”. **H.** Validation of endogenous expression pattern of IRAK4 (Left) and NEK1 (Right) in the regenerating neonatal heart. Positive signals are indicated by arrow. **I.** (Top) Schematic of IRAK4 expression cassettes in AAV6 constructs. Overexpress IRAK4 (green) in NRVM causes sarcomere (stained with α-actinin, red) disassembly (i) and pT445 adducin (stained with pT445, red) translocated from nucleus to cytoplasm (ii). Arrowheads indicate adducin is expressed in the nuclei in IRAK4 negative cells. Scale bar = 10 μm. **J.** (Top left) Schematic of IRAK4 expression cassettes in AAV9 constructs. (Top right) Schematic drawing of experimental design. (Bottom) AAV9-IRAK4-GFP was injected into CD1 pups at P1 and hearts were collected at P15 for sectioning and staining. IRAK4 overexpression caused strong sarcomere disassembly (i) in cardiomyocyte compared to littermate controls (ii).

### Identification of kinases that mediate adducin phosphorylation in cardiomyocytes

Given the critical role that T445/T480 phosphorylation plays in sarcomere disassembly, we sought to determine the mechanism of this phosphorylation event. To identify kinases that mediate phosphorylation of the membrane-associated sites T445 and T480, we performed a kinase-screening assay of 245 kinases for the phospho-sites T445 and T480. The initial screen identified seven potential kinases with relative high activity values compared to background for the phospho site T450 (Table S1, Fig. 5F left). Peptide of T480 only showed a significant response to kinase VRK1 (Fig.5F right). We then performed a secondary screen on the top hits outlined above using a HitConfirmation™ dose-response assay. These studies confirmed the activity of only three kinases namely NIMA related kinase 1 (NEK1), Interleukin 1 Receptor Associated Kinase 4 (IRAK4), and Doublecortin-like and CAM kinase-like 2 (DCAMKL2) that displayed concentration-dependent phosphorylation of the T445 phosphosite (Fig. 5G). These kinases also showed a dose response increase on T480 as well, although this appeared less prominent than that of the peptide T445. Endogenous staining of kinase candidates on P4 heart confirmed that both IRAK4 and NEK1 showed clear colocalization with disassembled sarcomeres (Fig. 5H).

To determine the role of IRAK4 in induction of sarcomere disassembly, we generated IRAK4-AAV6 constructs to overexpress IRAK4 in neonatal cardiomyocytes in vitro. Intriguingly, overexpression of IRAK4 resulted in sarcomere disassembly (Fig. 5I, Top panel), which was associated with expression of phosphorylated adducin pT445 in the cytoplasm of IRAK4 positive cells (Bottom panel). In IRAK4 negative cells, adducin was only expressed in the nucleus (Fig. 5I, Bottom panel with arrowheads). NEK1 overexpression in NRVM did not have any clear effect on sarcomeres or adducin phosphorylation (data not shown).

To evaluate the sarcomere structural changes upon IRAK4 overexpression *in vivo,* we generated an AAV9-IRAK4 virus. We injected this virus into neonatal mouse hearts at P1 and collected hearts at P15, a timepoint well beyond cell cycle exit of neonatal cardiomyocytes. Immunostaining on heart sections demonstrated that cells with IRAK4 overexpression displayed prominent sarcomere disassembly (Fig. 5J). These results indicate that IRAK4 phosphorylates adducin in vitro and in vivo and induces sarcomere disassembly. Collectively, our results identify adducin heterodimer as an important regulator for cardiomyocyte sarcomere disassembly during cardiomyocyte mitosis

### Identification of binding partners of adducin complex in regenerative and non-regenerative timepoints

Our results thus far suggest that the phosphorylated adducin heterodimer mediates cardiomyocyte sarcomere disassembly. However, given that adducins have not been previously studies in contractile cells, the binding partners of adducin in cardiomyocytes are unknown. In order to identify proteins that interact with adducin in cardiomyocytes, we performed MS analysis on P1 and P7 heart protein extracts using ADD1 antibody as the bait (strategy similar to Figure 1A but using an ADD1 antibody instead of Tnnt2 antibody). We found that numerous targets that were identified from the initial Tnnt2 pulldown assay were also detected in α-adducin pulldown (Fig. 6A), suggesting that α-adducin associates with sarcomeric proteins in the early postnatal heart.

**Figure 6.**
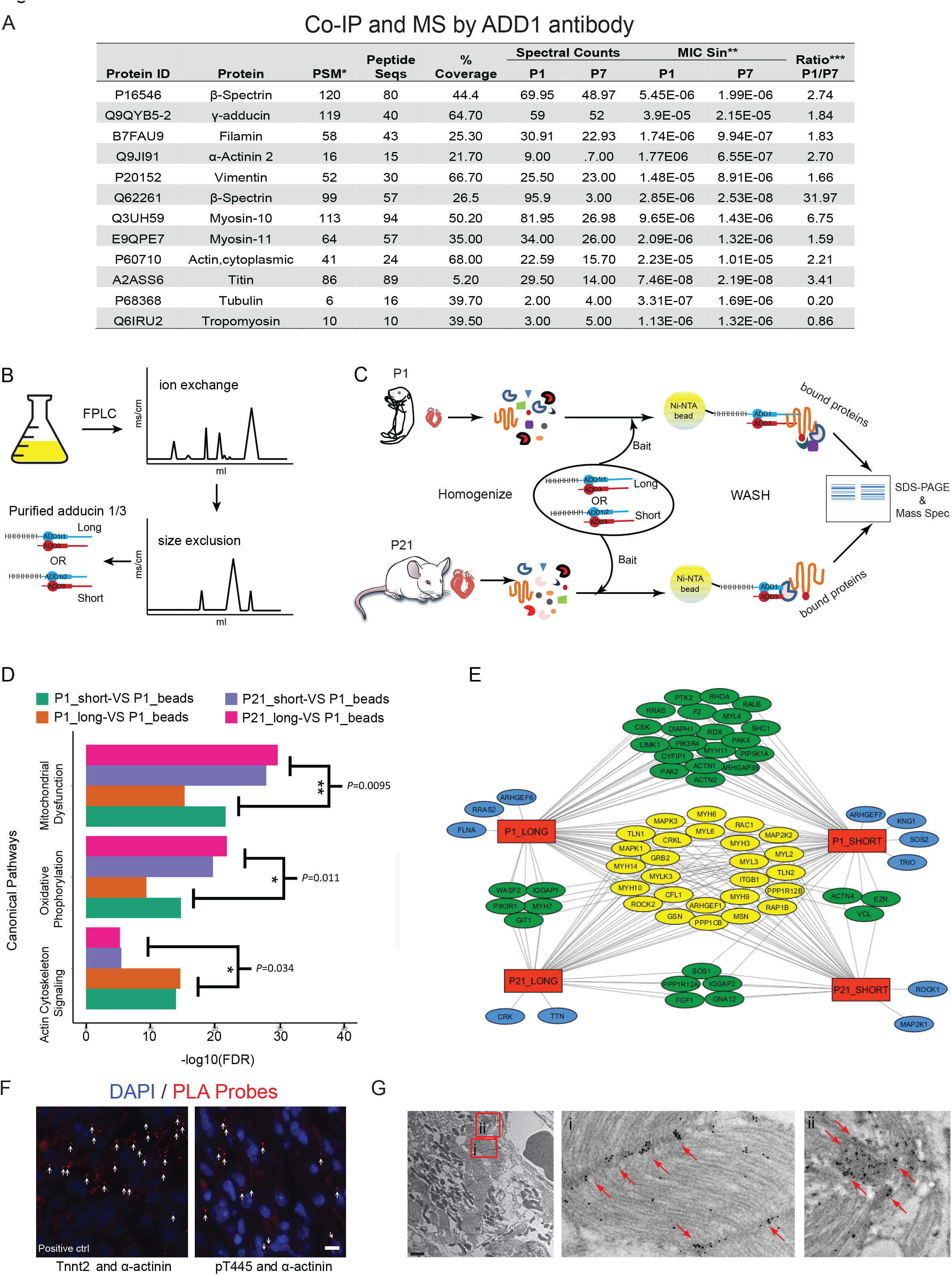
Analysis of Co-IP/MS with purified adducin complex. **A.** Table lists selected ADD1 associated proteins identified from the Co-IP/MS. * PSM: Peptide Spectrum Matches. Number of spectra assigned to peptides that contributed to the inference of the protein. **MIC Sin: The normalized spectral index statistic for the protein for the specific group. Calculated from the intensity of fragment ions in each spectrum assigned to a particular protein. ***Ratio: Quantitative ratio for protein between groups derived from the MIC Sin value. Scale bar=10 μm. **B**. Schematic of the purification process of short and long adducin complexes. **C.** Schematic drawing shows the flowchart of Co-IP/MS by purified adducin complex pulldown **D.** Bar graph for Canonical Pathways enriched in each sample. X-axis shows transformed adjusted *p* value (Benjamini-Hochberg Procedure) of overlap between sample proteins and pathways. FDR refers to false discovery rate. *P<0.05, **P<0.01, ***P<0.001; Statistical significance was calculated using a two-tailed *t*-test. **E.** Network plot of cytoskeleton proteins from the pull down assay interact with adducin_long or adducin_short at different time points (P1 and P21). Different color labeling highlighting common and unique proteins among sample groups. Red color represents the four sample groups in the pulldown assay. Yellow color represents proteins common to all groups. Green represents proteins common to only two or three groups. Blue represents proteins unique to each group. **F.** Proximity ligation assay (PLA) to examine direct interaction between adducin (pT445) and sarcomeric proteins (α-actinin 2) in P4 heart tissue. The red dots (arrows) are fluorescent signals which indicate a close interaction between two antigens. Samples include Interaction between α-actinin and troponin T (as positive control) or phospho-adducin (pT445) and α-actinin. Scale bar=10 μm. **G.** Immuno-EM image to show the subcellular location of phospho-adducin in cardiomyocytes with disassembled sarcomeres in neonatal MI hearts at day 3 after surgery. Two red boxes in left image are magnified on the right. (i) adducin is localized to z-disks. (ii) phospho-adducin is heavily localized to z-disks associated with plasma membrane in cardiomyocytes with disassembled sarcomeres. Red arrows indicate location of adducin.

Because adducin functions as a duplex (α/γ in cardiomyocytes), using ADD1 antibody as the bait might not recapitulate its binding pattern. In addition, while the aforementioned antibody-based pulldown might work in the neonatal heart, the lack of meaningful adducin expression in the adult heart precludes from conducting this study in adults. To further understand why even in the dTG hearts disassembly appears to be more pronounced in neonatal compared to adult cardiomyocytes, we conducted a pulldown study using purified α/γ-adducin complex as bait with either P1 vs P21 heart extracts. We cloned both Add1 and Add3 genes into the plasmid pETDuet which co-express two ORFs together and form a duplex naturally in cells. 6xHis-tag was cloned at the N-terminus of Add1 to facilitate purification and pulldown assay. To determine whether the long (ADD1i1/ADD3) and the short (ADD1 i2/ADD3) forms of adducin differentially bind sarcomeric proteins during the regenerative window compared to later timepoints, we generated two plasmids to co-express ADD1i1/ADD3 (long form) or ADD1i2/ADD3 (short form) respectively. Purified proteins were incubated with heart lysates overnight then bound to Ni-NTA beads to allow adducin and associated proteins attach to the resin. We also included a negative control where heart lysates were mixed with Ni-NTA beads. Figure 6 B and are schematics of the protein purification and Co-IP using the purified protein complex as bait. Samples and their corresponding controls were then analyzed by mass spec in form of log2(sample/control) to calculate fold change. Fold changes greater than 1.5 were considered for further Ingenuity Pathway Analysis. Fig.6D is a bar graph of statistically significant canonical pathways enriched in sample. Intriguingly, the only upregulated pathway at P1 compared to P21 is the actin cytoskeleton signaling pathway (two other mitochondrial-related pathways were downregulated at P1 compared to P21). We also generated a network plot to shows how cytoskeleton signaling pathway proteins interact with long or short adducins at different postnatal time points (P1 and P21) (Fig.6E). We also found that both the long and short forms of adducin exclusively bind specific sarcomeric proteins in P1 but not P21 hearts. Specifically, we found that the adducin complex binds cardiac actinin (ACTN2) only in P1 but not P21 hearts. In retrospect, this perhaps should not have been surprising given that actinins are scaffolding proteins which belong to the spectrin superfamily^44^, and spectrin is a known binding partner of adducins in non-contractile cells. In cardiomyocytes, alpha actinin is localized to the Z disks, where it is critical for regulating sarcomeric actin organization and sarcomere assembly. Therefore, a potential interaction between adducin and alpha actinin may be a mechanism of sarcomere disassembly during cardiomyocyte mitosis. To assess the interaction between adducin and α-actinin2, we performed proximity ligation assay (PLA) (Duolink®, Sigma-Aldrich, St. Louis, MO) (Fig.6F) using anti-phospho T445 antibody in the neonatal heart. For positive control, we used TNNT2 binding with α-actinin 2 in PLA assay. Or results demonstrate that adducin pT445 associates more with ACTN2 *in vivo* in the neonatal heart. In addition, we used Immunogold electron microscopy (Immuno-EM) imaging to determine the subcellular localization of pT445 adducin in neonatal cardiomyocytes with disassembled sarcomeres, which demonstrated that adducin is localized to the z-disks of sarcomeres (which is also where alpha actinin is localized) (Fig.6 Gi). pT445 adducin also appears to be associated with disassembled sarcomeres at the cellular cortex in association with the plasma membrane (Fig. 6Gii).

## Discussion

Numerous studies have identified mechanisms of cardiomyocyte cell cycle regulation in recent years. These mechanisms include diverse pathways including direct cell cycle regulators, transcription factors, micro RNAs, Lnc-RNAs as well as non-myocyte factors including extracellular matric, immunological, and environmental factors among others. However, striated muscle-specific mechanisms that regulate sarcomere organization during cardiomyocyte mitosis remain poorly understood. In the current report we demonstrate that the cytoskeleton protein adducin regulates sarcomere disassembly during cardiomyocyte mitosis. In non-contractile cells, previous studies have identified mechanisms that regulate cytoskeletal assembly and actin sheets dimerization. For example, the cytoskeletal regulatory protein adducin is a major regulator of actin bundling, and actin-spectrin interaction^29,30^. The adducin family is comprised of 3 members, namely α-, β-, or γ-adducins. The fully assembled adducin is comprised of heterodimers of homologous alpha and beta or gamma subunits. Despite the known role of adducin in actin assembly in non-contractile cells, its role in contractile cells is entirely unknown.

Our results stem from a proteomics screen of proteins that differentially associate with TNNT2 during an early regenerative timepoint of the newborn mouse heart. Although several cytoskeleton proteins appeared to be differentially associated with TNNT2 during this early time point, we focused on adducin given its known cytoskeleton regulatory role in non-contractile cells. We provide several lines of evidence that support the association of adducin with sarcomeric proteins; first, immunostaining demonstrates that α-adducin is localized to the cytoplasm and membrane only in cardiomyocytes with disassembled sarcomeres. Second, Co-IP using ADD1 antibody or purified α/γ-adducin complex suggests possible association of adducin with sarcomeric proteins, in particular α-actinin. Third, we identified a membrane-associated phospho-isoform of Add1, which binds α-actinin by proximity ligation assay, and colocalizes with membrane-associated sarcomeres by Immuno-electron microscopy labeled with nanogold particles. These results indicate that Add1 is a sarcomere associated cytoskeletal protein in cardiomyocytes.

Importantly, gain of function studies in vitro and in vivo to demonstrate that forced adducin expression induces sarcomere disassembly. We showed that expression of several isoforms of adducin induces sarcomere disassembly in primary neonatal cardiomyocytes in culture. In addition, both single TG Add1 i2 and Add1 i1 phoshpho-mimic manifest a non-homogeneous expression pattern which resulted in extension of the sarcomere disassembly window until P14, but not to adulthood. Protein stability assays demonstrated that co-expression of Add1 or Add3 is necessary for the stability of both proteins, and this was confirmed in vivo in the dTG lines which displayed stable expression through adulthood. We also demonstrate that although the non-phosphomimic dTG lines display persistent adducin expression, this did not result in pronounced sarcomere disassembly, while the same dTG construct with a constitutively active T445/T480 phosphomimic resulted in sarcomere disassembly that persisted to adulthood.

One important limitation of the current findings is the incomplete disassembly that we observed in adult cardiomyocytes even in the phospho-mimic dTG. Although we noted complete disassembly in the early postnatal period, most adult cardiomyocytes displayed partial disassembly that appeared to be localized to the zone surrounding nuclei and in the periphery of cardiomyocytes with complete clearance of the sarcomeres in these zones. This partial disassembly likely explains the lack of appreciable decline in LV systolic function in this TG line. The mechanism of this differential effect of adducin on early postnatal compared to adult sarcomeres is unclear but might be related to changes in sarcomere phenotype with age. To address this question, we used purified alpha (either short or long isoform)-gamma adducin protein complex to identify binding partners in early neonatal and adolescent hearts. The results demonstrate that the adducin complex differentially binds to cytoskeletal proteins depending on the postnatal age. Specifically, both short and long forms of the adducin complex appear to bind selectively to the cardiac-specific α-actinin (ACTN2) in P1 but not P21 hearts. PLA and immunogold-EM support the notion that adducin associates with α-actinin in z disks. Although adducin has not been previously shown to bind α-actinin, α-actinin belongs to the spectrin superfamily, and spectrin is a known binding partner of adducin in non-contractile cells. Given the known role of α-actinin in anchoring actin to z-disks in cardiomyocytes, future studies should explore the potential role of α-actinin in regulation of sarcomere disassembly downstream of adducin. Collectively, these results identify an important regulatory mechanism of sarcomere disassembly which links cytoskeleton organization to cell cycle regulation in mammalian cardiomyocytes and provide insights into the challenges of mitosis in actively contractile cells.

## METHODS

### Experimental Animals

All protocols were approved by the Institutional Animal Care and Use Committee of the University of Texas Southwestern Medical Center. Both male and female mice were used in all experiments to ensure age-match and gender-match. Healthy mice were chosen randomly from the expansion colony for all experiments. CD1 mice were used for wild-type studies histology staining and AAV9 injections. The number of animals used for each experiment was 3 to 5.

### Co-Immunoprecipitation (Co-IP) and Mass Spectroscopy (MS)

Mouse hearts from two different regenerative time points (n=3 for each), 3 days after myocardial infarction (MI) at P1 (P1MI), and 3 days after MI at P7 (P7MI) were harvested and lysed in IP lysis buffer: 2.5mM Tris, pH7.4, 150 mM NaCl, 1 mM EDTA, 1% NP40, 5% glycerol. Proteinase inhibitor cocktail and phosphatase inhibitor were included at all times. 5 μg TNNT2 mouse monoclonal antibody (Thermo Scientific) was used for binding with 1 mg digested heart samples with gentle rocking at 4 °C overnight. Protein A agarose beads (Calbiochem) was added to the mixture and incubation was continued with gentle rocking for 3 hrs at 4 °C. After microcentrifugation for 30 seconds at 4 °C, the pellets were washed five times with 1ml IP lysis buffer. Final pellets were resuspended with 25 ul 2XSDS sample buffer and loaded on SDS-PAGE gel (4-20%). When samples entered the resolving gel for about 5-10 mm, the run was stopped, and the gel was stained with Coomassie Blue. The stained area was cut and diced into 1mm cubes. The samples were then subjected to mass spectrometer analysis for protein identification. Co-IP with adducin antibody (H-10, cat# sc-25731 Santa Cruz) was performed using the same protocol.

### Histology and Immunostaining

Heart tissues were fixed in 4% paraformaldehyde (PFA)/PBS overnight at room temperature and then processed for either paraffin or cryo embedding. H&E staining were performed according to standard procedures at UTSW core histology facility on paraffin sections. For cryo embedding, fixed tissues were incubated in 30% sucrose/PBS at 4 °C until the tissues sunk. Tissues were embedded in freezing medium, frozen at −80°C and prepared for cryosection. Immunostaining was performed according to previous description^45^. Briefly, after antigen retrieval, sections were permeabilized and blocked with 10% serum /0.3% Triton-X100/PBS for 20 min at room temperature. Then samples were incubated overnight at 4°C with primary antibodies. Dilutions of primary antibody and secondary antibody (ThermoFisher) incubation varied depending on experiments. After washing with PBS, the sections were stained with DAPI (ThermoFisher D1306) and mounted with VECTASHIELD (Vector Laboratories H1700) for imaging.

### Antibodies

Primary antibodies: anti-troponin T, cardiac isoform Ab-1, clone 13-11 (Thermo scientific MS-295-P1, 1:100), anti-sarcomeric α-actinin (Abcam ab68167), anti-phospho Histone H3 Ser10 (Millipore 06-570, 1:100), anti-α-adducin antibody (1B1, Developmental Studies Hybridoma Bank, 1:50, for Figure 1F), anti-α-spectrin antibody (3A9, Developmental Studies Hybridoma Bank, 1:50, for Figure 1F), anti-Filamin 1 (E-3, sc-17749, Santa Cruz, 1:100, for Figure 1F), anti-α-adducin (phospho T445) antibody (Santa Cruz sc16738, for WB and staining, 1:100), anti-α-adducin antibody-N-terminal (Abcam ab151474, for Add1i2 WB and TG), anti-α-adducin antibody (Santa Cruz H-100 sc-25731, for WT mouse WB and staining on pTRE-Add1 T445/T480E transgenic, 1:100), anti-Y adducin antibody (Santa Cruz sc-365177, clone G-2, for WB and staining, 1:100), anti-α-adducin (phospho S726) antibody (Abcam ab53093, 1:100. Note that phosphositeS726 in human is located at 724 in mouse. Therefore, we used S724 refer to this site in the paper), anti-α-adducin phospho S355 antibody (Abmart 20592-1NB-3/C63-S, 1:100, for WB and double transgenics) anti-FLAG antibody (M2, F3165, Sigma-Aldrich 1:100). Myosin Light Chain 2 antibody (ptglab, 10906-1-AP, for staining), anti-MYH6 antibody (Sigma-Aldrich, HPA001349).

### Imaging

Fluorescent tissue or cell images were obtained with Leica DM2000, Nikon Eclipse Ni microscope, or Nikon A1R confocal microscope and processed with Photoshop CS3 to generate merge images with different colors. To count cardiomyocyte cell size, images of WGA stained samples were analyzed by Fiji software.

### Western Blotting

Western blot was performed using standard protocols. Antibodies used in WB are anti-α-adducin antibody (Santa Cruz H-100, sc-25731, 1:500), anti-γ-adducin antibody (Santa Cruz H-60, sc-25733, 1:500), anti-FLAG antibody (Genescript #A00187, 1:3000), anti-Ty1 antibody (Diagenode, #C15200054, 1:3000) and Peroxidase AffiniPure donkey anti-rabbit IgG (Jackson ImmunoResearch 711-035-152, 1:20,000), Peroxidase AffiniPure goat antimouse IgG (Jackson ImmunoResearch 115-035-003, 1:20,000) were processed by LI-COR.

### Neonatal Rat Ventricular myocytes (NRVM) isolation and cell culture

Cardiomyocytes were isolated from the left ventricle of 1–2-day-old Sprague-Dawley rats following the protocol from an isolation kit (Cellutron Life Technologies, cat#nc-6031). Myocytes were then plated on cover slips coated with laminin (Life tech #23017-015) at a density of 1250 cells/mm in DMEM : M199 (3:1) containing 10% horse serum, 5% FBS, and 1% Penicillin/Streptomycin. 100 μmol/L 5’-bromo-2’-deoxyuridine is included to inhibit growth of fibroblast. 24 hrs after cell attachment, change culture medium.

### Cardiomyocyte Isolation

The isolation of cardiomyocytes from adult mouse hearts has been described before (Mahmoud et al., 2013). Briefly, adult hearts were freshly harvested without atria. Samples were fixed in 4% PFA overnight at 4 °C. Scissors cut at aorta area for better penetration. After brief wash in PBS to remove PFA, the hearts were incubated with fresh collagenase D (2.4 mg/mL, Roche) and B (1.8 mg/mL, Roche) supplemented with 1% Penicillin/Streptomycin for 12 hours at 37 °C using end-over-end shaker. The supernatants were collected and stored at 4 °C. After several rounds of digestion until no obvious tissue, the supernatants were combined and filtered via 160 μm nylon mesh filter. Final cells were collected by centrifugation at 1000rpm for 5 mins. Cell pellets were resuspended with fresh PBS supplemented with 1% P/S. For nucleation counting, at least 300 cardiomyocytes per sample were quantified by confocal z-stack imaging.

### Plasmids and mutagenesis

Mouse Add1 cDNA transcript variant 1 was purchased from Origene (Cat# MR210357). The Add1 gene was amplified and subcloned into pUC19 for point mutations by the GeneArt Site-Directed Mutagenesis Plus Kit (Life Technologies, cat# A14604).The Add1 variant 2 was cloned from P1 mouse cDNA library Mouse Irak4 cDNA. was purchased from Origene (MR207322). Primers used to generate site-directed mutagenesis or other clones are listed in supplements.

### Production and purification of recombinant AAV vectors

The plasmids pTR-GNP for generating AAV expression cassette, pDP6rs and pDG9 for packaging AAV6 and AAV9 respectively, are gifts from Dr. Roger Hajjar (Mount Sinai School of Medicine, New York, NY). We used helper virus-free, two-plasmid-based AAV packaging system for viral production. The viruses were produced by polyethylenimine (Polysciences, Warrington, PA)-mediated transfection in HEK293T cells (Human embryonic kidney, ATCC, #CRL-11268, Manassas, VA). After transfection for 72 hrs, the cells were harvested by centrifugation. After three rounds of freeze and thaw, the cell lysates were treated with Pierce Universal Nuclease for Cell Lysis (Pierce, Waltham, MA). The crude viral suspensions were subjected to two rounds of iodixanol density gradient centrifugation. The final concentrated viruses were dialyzed against PBS and titrated by quantitative real-time PCR.

### Cell culture, transfection and cycloheximide treatment

HEK293T cells were transfected with p-3xFlag-Add1i1 and p-3xFlag-Add1i2 in the presence or absence of p-3xTy1-Add3 plasmid. 24 hours after transfection, cells were treated with 80 μM cycloheximide. Cells were collected 0, 24, and 48 hrs after treatment. Protein were extracted and measured concentration. 10 μg total proteins were separated by SDS-PAGE and later transferred onto nitrocellulose membrane for WB. Anti-FLAG antibody was used to detect α-adducin expression. Anti-Ty1 antibody was used to detect γ-adducin expression. GAPDH was used as internal control.

### Generation of transgenic mice

ORF of constitutively active form of Add1i1 (T445/T480E) was subcloned into a pTRE-Tight DNA vector to generate the pTRE-T445/T480E plasmid. pTRE-T445/T480E was linearized with *PvuI* and microinjected into fertilized oocytes to generated transgenic mice using standard procedures^17^. To induce Add1 overexpression in cardiomyocytes, pTRE-T445/T480E mice were crossed with αMHC-tTA mice. Age matched αMHC-tTA mice were used as controls. cDNA of Add1 i2 were synthesized directly from P1 CD-1 mouse heart. The ORF region was subcloned into pBluescript based vector which promoter was substituted with αMHC promoter for specific expression in cardiomyocytes^46^. Plasmid αMHC-Add1i2 was linearized with *EcoRV* and microinjected into fertilized oocytes to generated transgenic mice. The transgenic mice were crossed with C57B6N/J. cDNA of Add3 was purchased from Genecript (GenScript Bioteh, NJ). The ORF was subcloned into the same vector as that of the αMHC-Add1i2. To generate Add1i2/Add3 double transgenics, linearized DNA of αMHC-Add1i2 and αMHC-Add3 were both microinjected into fertilized oocytes using standard procedures. To generate phospho-adducin double transgenic, mutated Add1 i2 T445E/T480E were subcloned under the promoter of αMHC. Linearized DNA of αMHC-Add1i2 T445E/T480E and αMHC-Add3 were microinjected into fertilized oocytes. Both double transgenic and phospho-double transgenic were crossed with C57B6N/J for breeding strategy.

### Immuno-Electron Microscopy

To label adducin antibody with nanogold particles, we used α-adducin phospho-T445 antibody gold labeling. Labeling procedure followed the protocol by J.R.Thorpe^47^.

### Proximity Ligation Assay

Proximity ligation assay (Duolink, #DUO92101, Sigma) was performed according to instructions. Cyro-sections from 3 days post P1MI samples were used for all studies. Antigen retrieval and permeabilization steps were the same as per standard imunnostaining procedure. Primary antibodies TNNT2 and α-actinin were used as positive controls.

### TUNEL Assay

Cryo-sections were stained for troponin T as abovementioned “Histology and Immunostaining” part. Following incubation with corresponding secondary antibody conjugated to Alexa Fluor 555 (Invitrogen), TUNEL staining was performed according to manufacturer’s guideline (In-Situ Cell Death Detection Kit, Fluorescein, Roche). All staining was performed on 3 hearts per group, with 3 sections per heart.

### Serine/Threonine Screening Assay

We synthesized two biotinylated peptides with the phosphorylation sites in the middle of the sequences. They are for “T445”, KQQREK**T**RWLHSG, and “T480”, EDGHR**T**STSAVP (characters labeled in red are the phosphorylation sites). The kinase screening assay was performed by ProQinase GmbH (Freiburg, Germany). The initial screening assay was determined at 1 μM in singlicate in a radiometric assay (^33^PanQinase® Activity Assay) on a panel of 245 Ser/Thr kinases, using streptavidin-coated FlashPlate® PLUS plates (PerkinElmer, Boston, MA). The reaction cocktails were pipetted into 96 well, V-shaped polypropylene microtiter plates. Each assay plate contains one well which has no enzyme and is used as background. For evaluation of the results, the background signal of each kinase (w/o biotinylated peptide) was determined in parallel. For Hit Confirmation assay, seven kinases picked from the first screening assay were tested with sample peptides in triplicate at three concentrations (1, 0.5, 0.25 μM).

### Protein expression and purification

Full length mouse ADD1 isoform 1 and ADD3 or mouse ADD! Isoform 2 and ADD3 were subcloned into the first and second Open Reading Frame (ORF) of pETDuet vector, respectively. mADD1 has 6xHis tag at its N terminus followed by TEV protease recognition site and the plasmid was transformed into Rosetta (DE3) pLysS cells (Novagen). Target protein was expressed in cultures grown in autoinduction media at 18°C overnight ^48^. The culture was harvested and lysed through French Press in lysis buffer (20 mM Tris (pH8.0), 150 mM NaCl, 0.5 mM DTT and supplemented with protease inhibitors). The lysate was centrifuged and the supernatant was load onto a Ni-NTA affinity column (Qiagen). Proteins were eluted with elution buffer (20 mM Tris (pH 8.0), 150 mM NaCl, 2 mM DTT and 250 mM Imidazole (pH 8.0)). The eluate was further purified by ion exchange chromatography (HiTrapSP) followed by gel filtration chromatography (GF_Superose6). The peak fractions were collected and concentrated to 2-3 mg/mL for later experiments.

### Co-IP of purified adducin 1/3

Purified adducin long form duplex ADD1 i1/ADD3 and short form duplex ADD1 i2/ADD3 are used as bait in the pull-down assay. Fresh P1 and P21 hearts were lysed in IP buffer: 50 mM NaH2PO4, 150 mM NaCl, 10 mM Imidazole, 0.05% Tween 20, pH8.0. Proteinase inhibitor cocktail and phosphatase inhibitor were included at al times. 2 μg purified adducin duplex (short or long) was used to bind with 800 μg digested heart samples with gentle rocking at 4 °C overnight. Ni-NTA agarose beads was washed 3 times with lysis buffer. Then the beads were added to the mixture. Incubation was continued with gentle rocking for 1 hrs at 4 °C. After microcentrifugation for 30 seconds at 4 °C, the pellets were washed with 1ml IP lysis buffer for 5 times. The IP lysis buffer was supplemented with 5mM Immidazole in the first 3 times; with 20 mM imidazole for the last 2 times. Ni-NTA beads bind with P1 or P21 heart lysates at 4 °C 1hr were included as a negative control. Final pellets were resuspended with 25 ul 2XSDS sample buffer and loaded on SDS-PAGE gel (4-20%). When samples entered the resolving gel for about 5-10 mm, the run was stopped and the gel was stained with Coomassie Blue. The stained area was cut and diced into 1mm cubes. The samples are then subjected to MS analysis for protein identification.

### Proteomics analysis

Abundances from Proteome Discoverer 2.4 were used to calculate fold changes (log2) between samples and their relevant controls. Only samples with an increase in fold change were considered for further Ingenuity Pathway Analysis (QIAGEN Inc., https://www.qiagenbioinformatics.com/products/ingenuity-pathway-analysis).

## Data analysis

All graphs represent average values, and all error bars represents standard error. All data collected and analyzed were assumed to be distributed normally. Unless specified, the two-tailed unpaired student’s *t*-test was used to determine statistical significance. \**P*<0.05, \*\**P*<0.01, \*\*\**P*<0.001 were considered statistically significant.

## Acknowledgements

We thank McDermott Center Sequencing and Bioinformatics Cores for sequencing and analysis, J. Richardson and J. Shelton for assistance with histology, H. Mirzaei, D.C. Trudgian, and A. Lemoff for performing MS and analysis, K. Luby-Phelps and A. Darehshouri for assistance with electron microscopy (1S10OD021685-01A1 to Katherine Luby-Phelps), R.E. Hammer, J. Ritter, M.Nguyen, H. Zhu for microinjections.

## Supplemental Figures

**Fig. S1.**
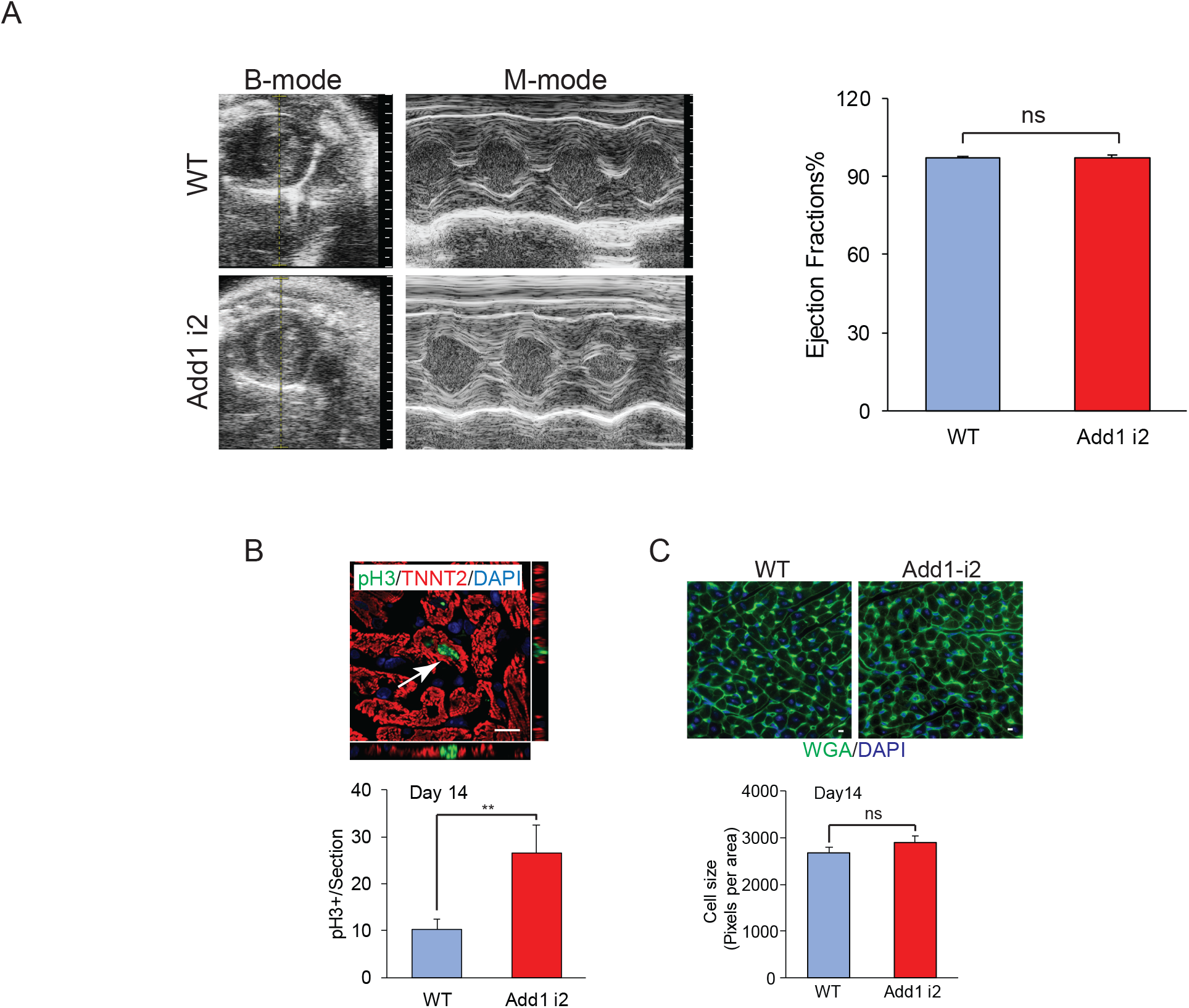
Characterization of adducin isoform 2 single transgenic. **A.** Echocardiography (Left) and left ventricular systolic function quantification by ejection fraction (Right) in 21 days ADD1 i2 transgenic. (n=3 for each group). **B.** (Top) Z-stack fluorescent image to show staining of pH3 (green) and TNNT2 (red) in P14 ADD1 i2 transgenic. (bottom) Graph to show the percentage of mitotic cardiomyocytes (arrow). (n=3 per group); Scale bar= 10 μm. **C.** (Top) WGA (green) and DAPI (blue) staining to show cardiomyocyte cell size. (bottom) Quantification of cell size by in ADD1 i2 transgenic. Cell measurements were processed by Fiji. Scale bar= 10 μm. Data are presented as mean ± s.e.m. *P<0.05, **P<0.01, ***P<0.001; statistical significance was calculated using a two-tailed *t*-test.

**Fig. S2.**
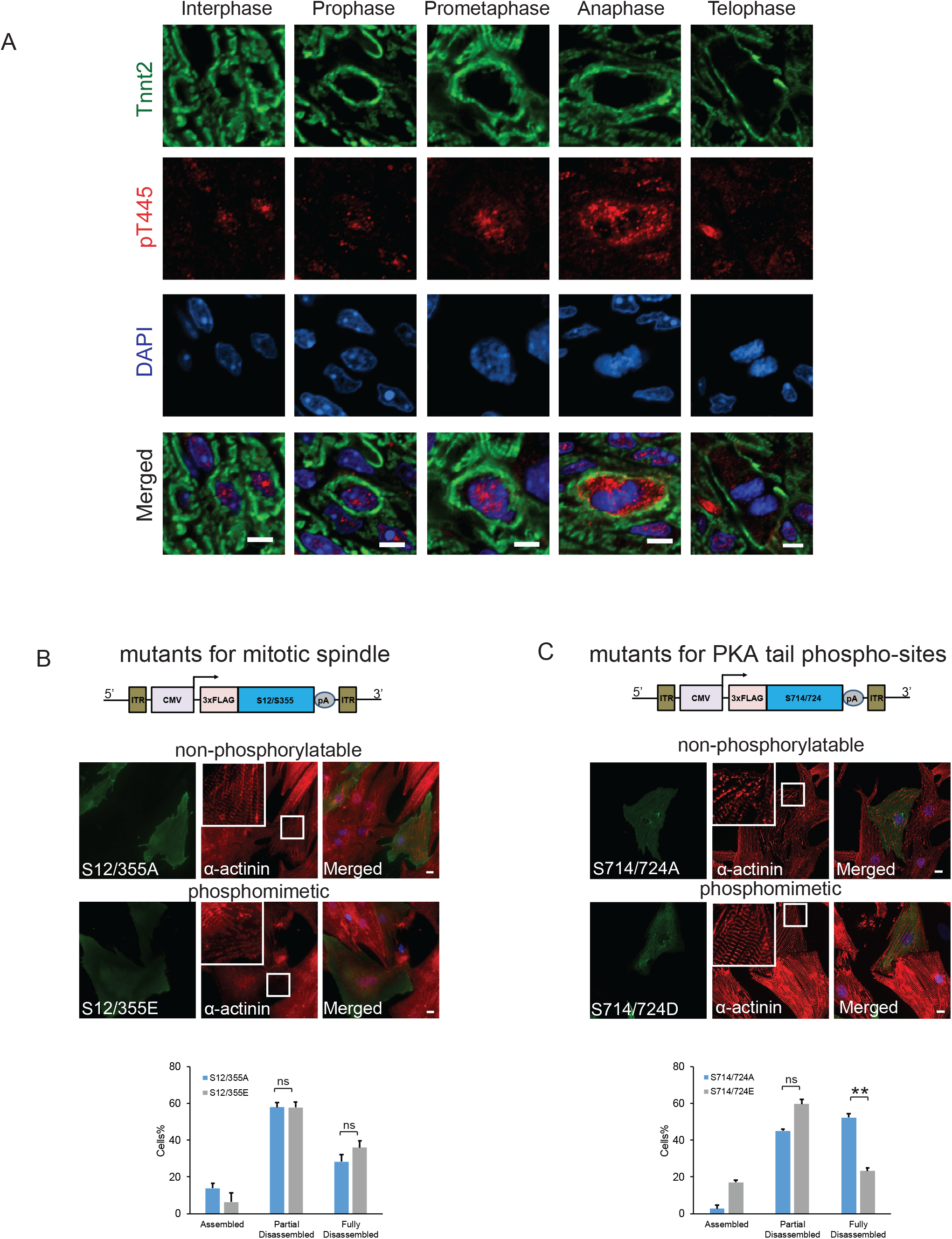
Effect of adducin phosphorylation on sarcomere disassembly. **A.** Phospho-adducin expression profile in regenerating neonatal heart at P4 by anti-pT445 staining. Note that the expression of pT445 translocates from nucleus to cytoplasm during cardiomyocyte mitosis. Scale bar= 10 μm. **B & C.** Overexpression of adducin mutants S12/S355 (**B**), and S714/724 (**C**) in NRVM induced by AAV6. (Top) Schematic diagram of AAV shuttle vectors for adducin expression. Mutants are expressed with 3 tandem FLAG epitopes in frame at the N-terminus. Middle. Immunofluorescent staining with α-actinin antibody (red) to show sarcomeric structure. Adducin mutants are detected by Flag antibody in green. *Insets* show high magnification images of boxed region. Bottom. Quantitative analysis of disassembled sarcomeres induced by adducin overexpression. Data are presented as mean ± s.e.m. *P<0.05, **P<0.01, ***P<0.001; statistical significance was calculated using a two-tailed *t*-test.

**Fig. S3.**
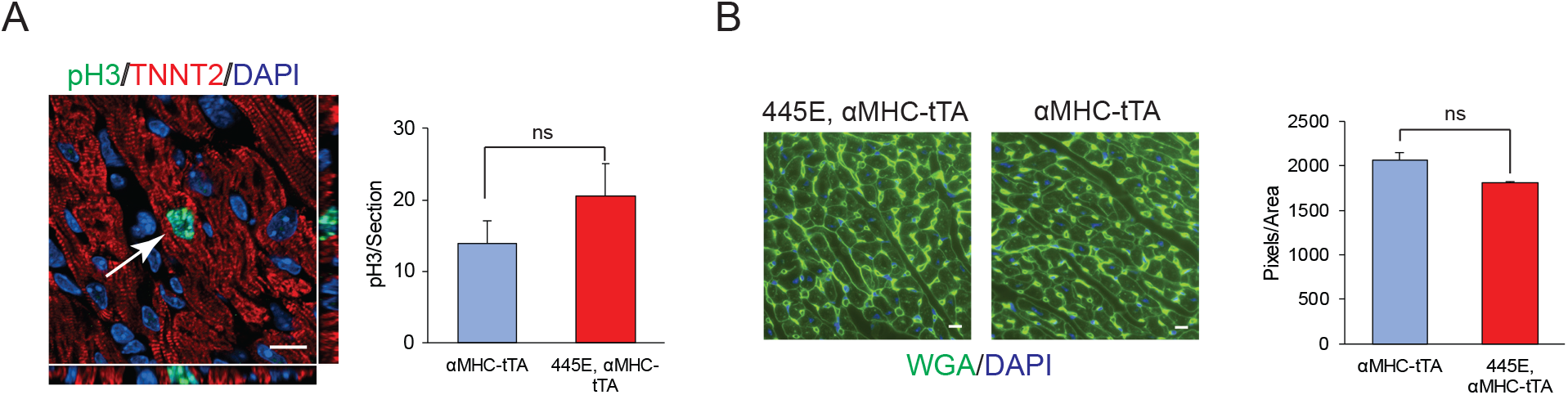
A. Characterization of cardiac inducible adducin phospho-mimic single transgenic. (Left) Z-stack fluorescent image to show staining of pH3 (green) and TNNT2 (red) in P14 inducible p-single TG transgenic. (Right) Graph to show the percentage of mitotic cardiomyocytes (arrow). (n=3 per group); Scale bar= 10 μm. **B.** (Left) WGA staining to show cardiomyocyte cell size. (Right) Quantification of cell size of the pTRE-Add1 transgenic. Cell measurements were processed by Fiji. Scale bar= 10 μm. Data are presented as mean ± s.e.m. *P<0.05, **P<0.01, ***P<0.001; statistical significance was calculated using a two-tailed *t*-test. Data are presented as mean ± s.e.m. *P<0.05, **P<0.01, ***P<0.001; statistical significance was calculated using a two-tailed *t-*test.

**Fig. S4.**
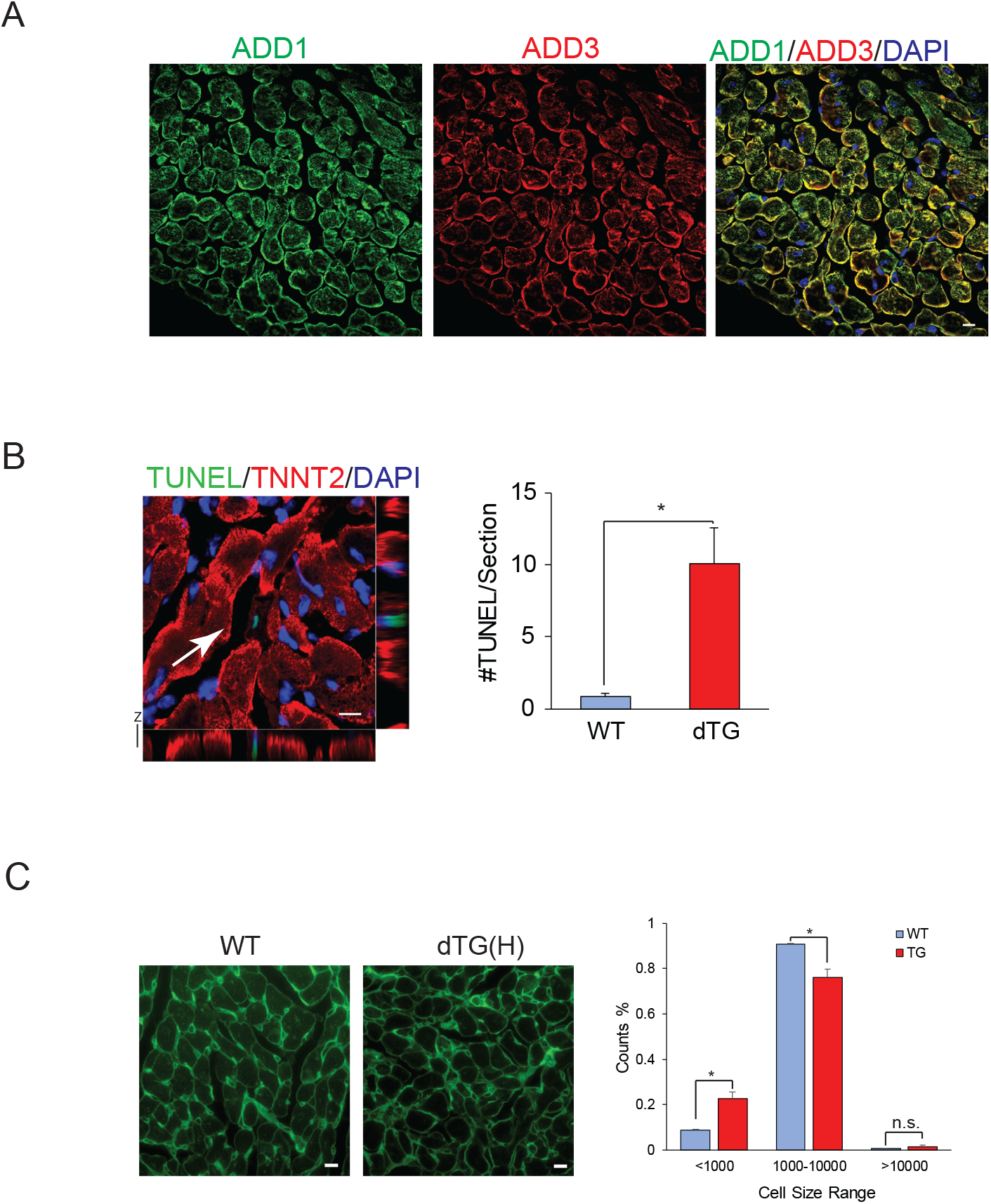
Characterization of cardiac-specific Add1i2/Add3 double transgenic. **A.** Co-stain of ADD1 (green) and ADD3 (red) in ADD1i2/ADD3 double transgenic to confirm colocalization in cardiomyocytes. Scale bar=10 μm. **B.** (Left) Immunofluorescence staining for TUNEL (green) and cardiac troponin T (red) in1-month-old ADD1 i2/ADD3 double transgenic. Arrow points to TUNEL positive cardiomyocyte. (Right) Quantification of positive signals per section. n=3 for each group. Scale bar = 10 μm **C.** (Left) WGA staining to show cardiomyocyte cell size. (Right) Quantification of cell size of WT and ADD1 i2/ADD3 double transgenic. Cell measurements were processed by Fiji. Because the double transgenic has many small cells, therefore, we categorize cell size into three counting groups. Scale bar= 10 μm. Data are presented as mean ± s.e.m. *P<0.05, **P<0.01, ***P<0.001; statistical significance was calculated using a two-tailed *t*-test.

**Fig. S5.**
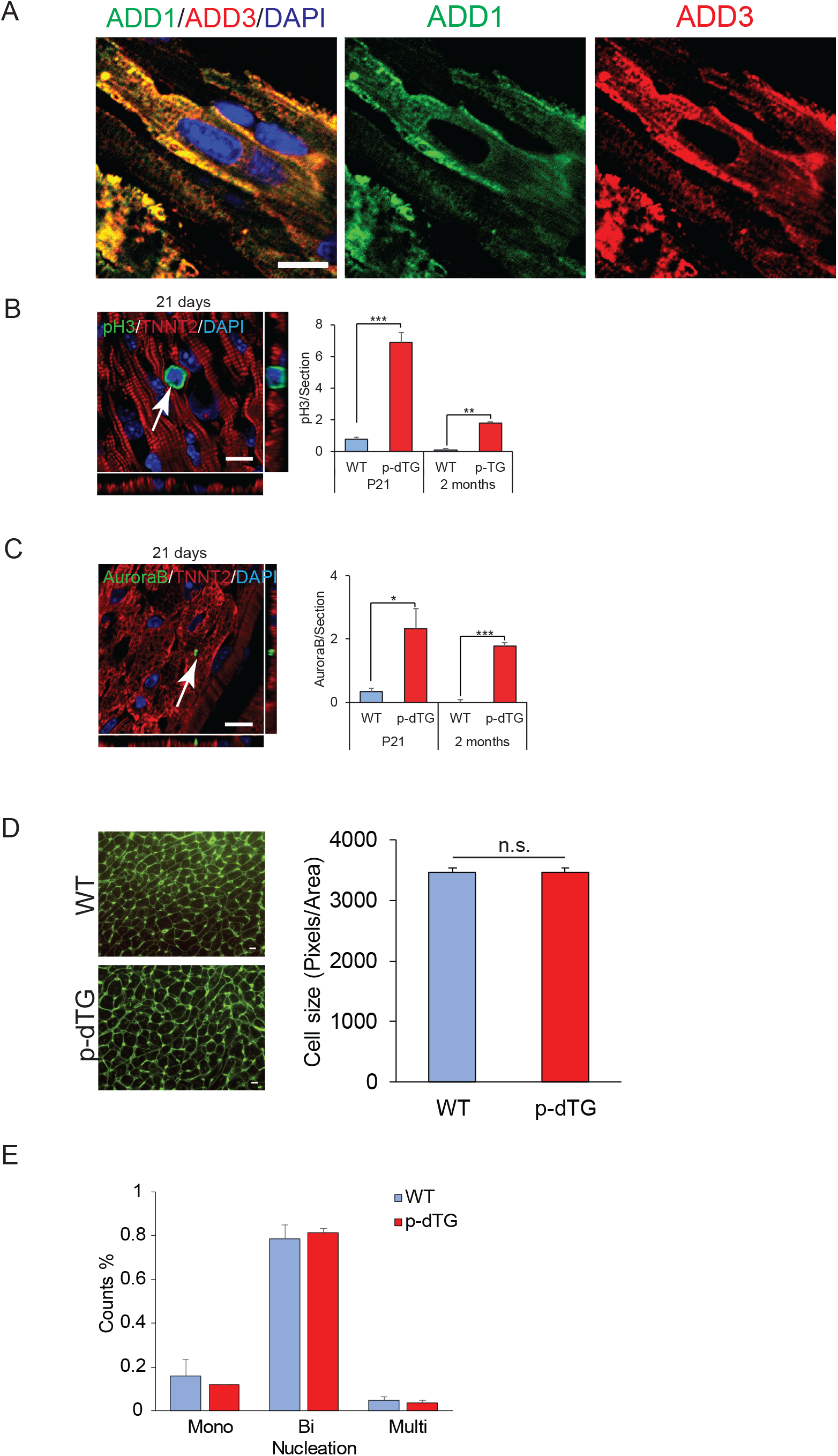
Characterization of cardiac-specific phospho double transgenic. **A.** Images to show colocalization of ADD1 and ADD3 in phospho-double transgenic. Scale bar= 10 μm. **B.** (Left) Z-stack fluorescent image to show staining of pH3 (green) and TNNT2 (red) in 21 days old phospho-double transgenic. (Right) Quantification of pH3 (green) positive cardiomyocytes (red) per section of P21 and 2 months hearts. (n=3 per group); Scale bar= 10 μm. **C.** (Left) Immunofluorescence staining for Aurora B kinase (green) and TNNT2 (red) of 21 days old p-dTG. (Right) Quantification of cytokinesis positive cardiomyocytes of both 21 days and 2 months old heart. n=3 per group. Scale bar = 10μm. **D.** (Left) WGA staining to show cardiomyocyte cell size. (Right) Quantification of cell size of WT and phospho-ADD1 i2/ADD3 double transgenic. **E.** Nucleation quantification of 2-month-old WT and phospho-double transgenic. (n=3 per group). Scale bar = 10μm.

---

**Supplemental Table 1:**

Raw data of assessment of the phosphorylation profile of peptide “T445” and “T480”at one concentration (1 μM) in 245 Ser/Thr kinase assay.

